# Automated derivation of mean field models from spiking neural networks for the simulation of brain dynamics

**DOI:** 10.64898/2026.03.18.712631

**Authors:** Roberta M. Lorenzi, Marialaura De Grazia, Claudia A.M. Gandini Wheeler Kingshott, Fulvia Palesi, Egidio D’Angelo, Claudia Casellato

## Abstract

A mean field model (MFM) is a mesoscopic description of neuronal population dynamics that can reduce the complexity of neural microcircuits into equations preserving key functional properties. The generation of a MFM is a complex mathematical process that starts with the incorporation of single neuron input/output relationships and local connectivity. Once neuron electroresponsiveness and synaptic properties are defined, in principle, the process can be automatized. Here we develop a tool for automatic MFM derivation from biophysically grounded spiking networks (Auto-MFM) by performing micro-to-mesoscale parameter remapping, estimating input/output relationships specific for different neuronal populations (i.e., transfer functions), and optimizing transfer function parameters. Auto-MFM was tested using a spiking cerebellar circuit as a generative model. The cerebellar MFM derived with Auto-MFM accurately reproduced cerebellar population dynamics of the corresponding spiking network, matching mean and time-varying firing rates across a wide range of stimulation patterns. Auto-MFM allowed us to model and explore physiological and pathological circuit variants; indeed, it was used to map ataxia-related structural connectivity alterations of the cerebellar network, in which Purkinje cells with simplified dendritic structure altered the cerebellar connectivity. Furthermore, Auto-MFM was used to create a library of cerebellar MFMs by sweeping the level of the excitatory conductance at mossy fiber - granule cell synapse, which is altered in several neuropathologies. Auto-MFM is thus proving a flexible and powerful tool to generate region-specific MFMs of healthy and pathological brain networks to be embedded in brain digital models.

## 1. Introduction

Mean field models (MFM) are a mathematical representation of mesoscopic dynamics, obtained by simplifying complex systems through the approximation of multiple elements interactions with an average interaction field^1,2^, by reducing a many-body problem into a one-body problem. Thanks to its nature, the mean field theory finds a broad range of applicability, spanning from physics to economics and neuroscience^3–5^. Applied to neuroscience, and more specifically to brain modeling, the mean field approximation enables the description of neuronal populations providing a statistical summary of interactions between thousands of neurons embedded in a microcircuit^6^. Although the mean field approach neglects the spatial organization of neuronal microcircuits, it reduces the computational burden of simulating neuronal systems providing effective representations of functional regions in large-scale brain models, where the explicit simulation of single-element microcircuits remains computationally intractable. Furthermore, experimental signals are typically acquired at the mesoscale level. Voxels in functional magnetic resonance imaging (fMRI) acquisitions or source fields in electroencephalography reflect the average activity of large neuronal ensembles, thus setting at a resolution compatible with the assumptions underlying mean field modeling^7^

Different mathematical approaches allow to derive MFMs depending on the microcircuit properties that need to be translated into the mesoscopic domain. Indeed, microcircuits can include neuronal models with different granularity and complexity. Detailed biophysical models, e.g., Hodgkin–Huxley style, include multiple compartments to represent the neuronal morphology and explicitly simulate ionic conductances and gating variables, offering the greatest biological realism at the cost of computational tractability^8^. Single-compartment models, i.e., point-neurons, overlook morphological details yet retaining the salient functional properties of neurons, thereby providing an efficient trade-off between physiological accuracy and computational tractability^9^. Commonly used representations of point-neurons include the Leaky Integrate-and-Fire (LIF) neuron, which represents membrane potential dynamics as a linear response to input current^10^, the Extended Generalized Leaky Integrate-and-Fire (E-GLIF) model, which features non-linear second order dynamics allowing for a flexible parametrization of different neuronal types^11^, the Adaptive Exponential Integrate-and-Fire (AdEx) model, which incorporates spike-frequency adaptation and nonlinear action potential initiation^12^, and the Quadratic Integrate-and-Fire (QIF) model, which enables analytical reductions due to its quadratic voltage dynamics^13^.

Multiple MFM derivation approaches can be exploited. Analytical mean-field approximations ^14–17^ yield closed-form solutions relying on a Lorentzian ansatz for the membrane potential distribution and are strictly constrained to the QIF model. Similarly, a biophysically-inspired MFM has been derived based on a network of Hodgkin–Huxley neurons, retaining ionic dynamics driving neuronal activity simulations^18^. Despite its high biophysical interpretability, its derivation depends on assuming a Lorentzian voltage distribution, as in the case of QIF-based MFM, which limits their validity to near-synchronous regimes. MFMs can also be derived via a master equation formalism, not assuming any specific voltage distribution and can be systematically applied to a broad class of neuron models including the ones that are not analytically tractable, thus providing a more general and flexible framework for constructing mesoscopic models from spiking neural network (SNN) simulations^19^. This formulation assumes that network dynamics are Markovian and that the transfer function (TF) captures single-neuron responses at the population level. The TF, in general, is the mathematical construct that allows to translate microscale properties into MFM equations.

A TF can account for multiple distinct neuronal populations, and its derivation requires a SNN, in which thousands of single-point neurons are connected^20,21^. Single-neuron parameters and the SNN activity provide priors for deriving the population-specific TFs. These are defined by three connection-specific parameters (for each pair of pre and post synaptic populations), namely the synaptic convergence, the quantal synaptic conductance, and the synaptic time decay. This procedure has been widely adopted in neuroscience, becoming the technical reference of biologically-grounded MFMs incorporating specific features of multiple neuronal populations in cortical and subcortical regions^22–26^.

Despite its broad applicability, the generation of these MFMs is a complex mathematical process that starts with the incorporation of single neuron input/output relations and synaptic features. When SNNs are constructed as biophysical representations of circuits, they are scaffolds that replicate in-silico the anatomical spatial constraints, the cell-specific morphology and orientation-dependent connectivity; the point-neuron parameters for each neuronal population are tuned on the corresponding multicompartmental models. Translating these properties into a mesoscale TF-based formalism is not straightforward. Here, we developed a procedure for the automatized generation of MFMs (i.e., Auto-MFM), with a modular architecture enabling automatized parameter extraction from SNN, considering the synchronization of synaptic transmission, and implementing a TF optimization to improve the match between input/output relation of SNN and MFM. We tested it on the cerebellar biophysical SNN, significantly improving the previous version based on trial-and-error tuning of TF gains^25^. To date, the cerebellar MFM is the most complex amongst the cortical and subcortical MFMs: it is made of four neuronal populations, comprising granule cells (GrC), Golgi cells (GoC), Stellate and basket cells, and Purkinje cells (PC), embedded in a multi-layer architecture with an input layer (i.e., granular layer including GrC and GoC), an hidden layer made of stellate and basket cells, and an output layer made of PCs^27^. Here, the cerebellar MFM was customized to reproduce population activities and mutual interactions characterizing in *vivo* states. Auto-MFM was then used to map the alterations of cerebellar structure and function of ataxic mice, in which dendritic simplification in Purkinje cells brings about altered connectivity and dynamics in the whole cerebellar SNN. Moreover, Auto-MFM was used to investigate alterations in GrC dynamics induced by changes in the glutamatergic input from mossy fibers (mfs), demonstrating the flexibility of the toolkit in generating a MFM library with specific parameter sweeps.

## 2. Methods

The auto-MFM tool has been developed as a framework to transparently translate microscopic properties into mesoscopic domain through the derivation of population-specific TFs (see the pipeline diagram in Fig. 1). Auto-MFM is a python-based open-source software publicly available on GitHub for the construction of region-specific MFMs, generalizable to multi-population single-point networks into a structured circuit volume. It can be part of the ecosystem around the Brain Scaffold Builder (BSB) component-based framework (https://www.ebrains.eu/tools/bsb)^28^,since it directly reads the configuration file of any brain region SNN and the spiking activity files from simulations in BSB-NEST (Fig. 1). Auto-MFM indeed includes an ensemble of functions to implement the automatized MFM creation from the “equivalent” SNN described in this section (Fig. 2): (i) the parameter-efficient transfer from SNN to MFM (section 2.1; Fig. 2A), including the description of single neuron models and biophysical SNN, along with the automatic refinement strategies for translating single-neuron spike events into population-mean signals ; (ii) the MFM derivation process (section 2.2; Fig. 2B), based on the master equation formalism to derive neuronal population-specific TFs; (iii) the optimization of population-specific TFs parameters (section 2.2; Fig. 2C). The application of this pipeline is described for the derivation of the cerebellar MFM (section 2.3), and the derivation of cerebellar circuit pathological variants (section 2.4).

**Figure 1.**
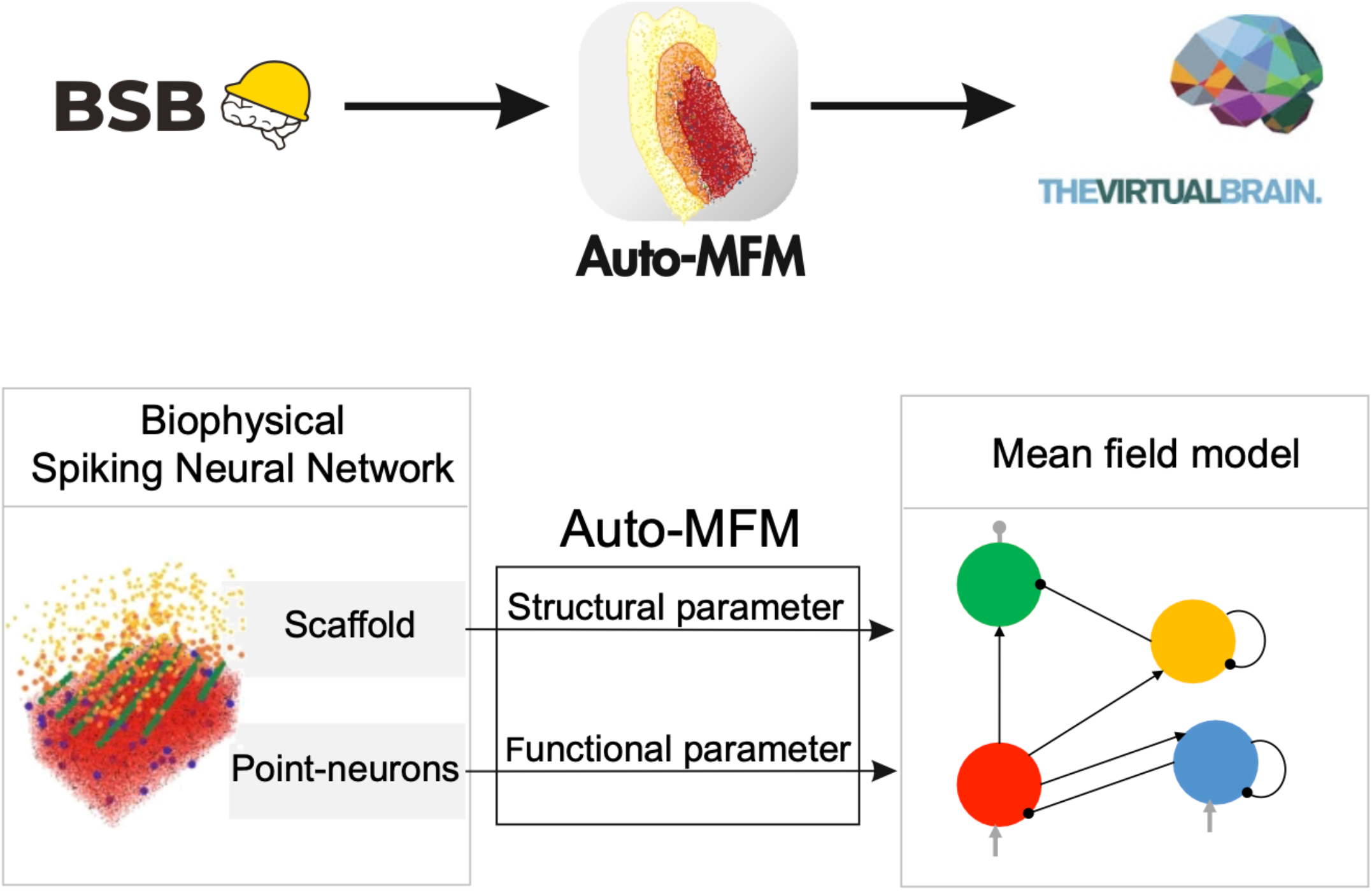
The multiscale brain modeling ecosystem. BSB (Brain scaffold builder: https://www.ebrains.eu/tools/bsb) with its interface with NEST is used to construct SNN with morphology-based connectivity rules, representing the microcircuitry of a brain region. BSB-NEST interface is used to simulate the point-neurons spiking activity embedded in the SNN. Auto-MFM (Automatic mean field derivation tool: https://github.com/RobertaMLo/Auto-MFM) is used to derive mesoscale modeling directly from BSB-based reconstruction. The auto-MFM toolkit executes a systematic extraction and resampling of the structural and functional parameters of biophysical spiking neural networks, generating mesoscale models that preserve spatial organization, neuronal population resolution, and signal propagation properties of the underlying microcircuit. TVB (The Virtual Brain: https://www.thevirtualbrain.org/tvb/zwei/) is used to simulate whole brain dynamics by representing brain region with computational model. MFM models integrated into TVB become region-specific functional units of the large-scale brain simulator.

**Figure 2.**
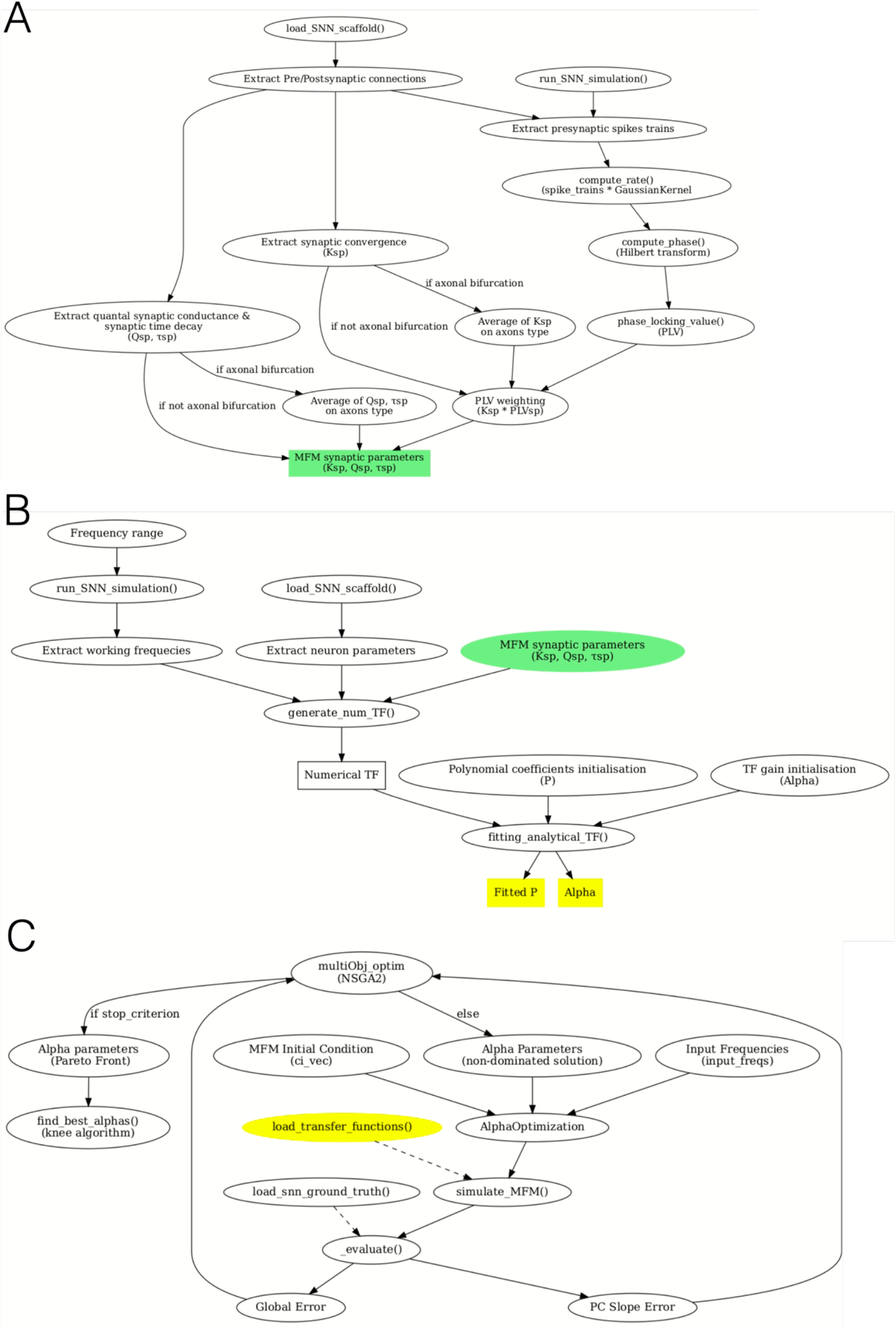
Auto-MFM modules. Colored square boxes are the output of each module. **A)** Parameter-efficient transfer from micro to mesoscale accounting for the structural details of the SNN, and the synchronization of incoming synaptic input quantified through the Phase Locking Value (PLV) of SNN oscillations; **B)** Derivation of the MFM through a population-specific TF fitting, using the synaptic parameters derived in A) (MFM synaptic parameters - in yellow). The procedure consists in the derivation of a numerical template from SNN (i.e., numerical transfer function), that is the reference for the computation of the analytical TF by fitting its polynomial coefficient (P) and tuning a computational gain (alpha - α) to fit both low and high frequency (Lorenzi et al., 2023). **C)** Optimization of the MFM, using genetic algorithm (NSGA2 – pymoo implementation) for a multi-objective optimization, targeting the specific α for each population. Population TFs computed in B) are loaded (yellow box) and used to generate the MFM activity. In the cerebellar test-bench, the objectives are the global error (i.e., the sum of population-specific errors across multiple input stimulations) and the output population slope and are optimized against the SNN reference (snn_ground_truth).

### 2.1 Parameter-transfer from micro to mesoscale

The basis for a derivation of a mesoscale model is the microscale computational representation. For the MFM derived via master equation formalism, microscale level is represented by SNN. However, when transitioning to MFMs, spatial information is not explicitly represented, therefore the first module of the auto-MFM tool is intended to properly map the parameters.

#### 2.1.1 Spiking Neural Networks (SNN)

The building blocks of SNN are the neurons represented by point-neuron models. Each neuron type is characterized by parameters tuned on the most salient features extracted from the electroresponsiveness of the corresponding multicompartmental neuron model and/or on experimental electrophysiological recordings and morphology digital reconstruction. The point-neuron model used here for the cerebellar MFM test-bench is the E-GLIF described by the following equation system^29,30^:

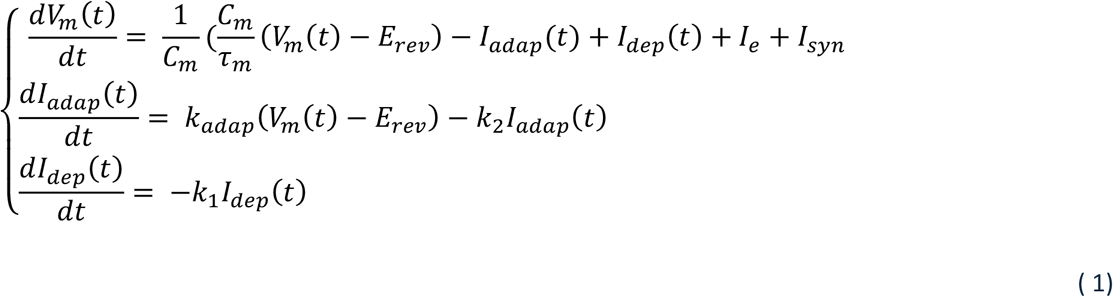

where V_m_ = membrane potential (mV), C_m_ = membrane capacitance (pF); τ_m_ = membrane time constant (ms); E_rev_ = reversal potential (mV); I_adap_ = adaptation current (pA); I_dep_ = depolarization current (pA); I_e_ = endogenous current (pA); I_syn_ = synaptic current (pA); k_adap_ and k_2_ = adaptation constants; k_1_ = decay rate of I_dep_.

The synapses between point-neurons are modelled as conductance-based, with the synaptic current (I_syn_) defined as:

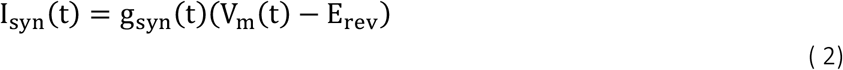

With g_syn_ (t) = conductance dynamics modeled as time-dependent kernels such as exponential or alpha functions.

Biophysical SNNs are constructed using connection rules retaining the anatomical organization of the microcircuit, typically at micrometer resolution^28^. When transitioning to MFMs, spatial information of the underlying biological circuit is not explicitly represented in the equations, thus requiring an overall mesoscale definition of the synaptic parameters that is implemented accounting for the presynaptic activity synchronization and axonal divergences.

#### 2.1.2 Temporal coupling of synaptic transmission

A biophysical SNN includes the parametrization of the structural synaptic densities in terms of synaptic convergences, i.e., the parameter K, for each connection type. Structural convergence induces an effective temporal coupling between the presynaptic neuronal activities. MFMs treat neurons of the same type as a homogenous population, and therefore, the value of K should be pruned considering the temporal coordination among the neurons of the same types converging on the same postsynaptic neuron. The phase locking value (PLV) was computed as the temporal coupling useful to scale down the structural convergence to an effective synaptic signal convergence. PLV was then used as weighting factor for K for each connection type, yielding a K that reflects both anatomical density and timing coordination for synaptic transmission.

To estimate the PLV, the SNN was simulated under a homogeneous Poisson signal for a fixed frequency (usually the half frequency of the range of interest) in the input fibers. For each connection type, a postsynaptic target neuron was chosen as representative of the population, and its presynaptic neurons were identified from the SNN connectome. From each presynaptic spike train, the firing rate signal was derived, and the analytic signal was computed via the Hilbert transform to get the phase time series. The PLV was calculated pairwise across all presynaptic signals, using ELEPHANT toolbox (doi:10.5281/zenodo.1186602; RRID:SCR_003833):

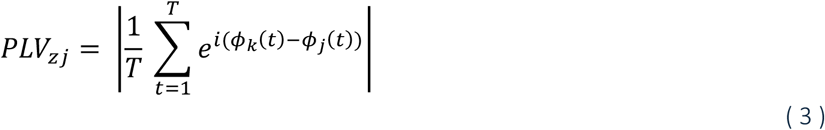

The PLV is computed between pairs of presynaptic neurons [*z,j*] projecting to the same postsynaptic neuron. *ϕ* denotes the instantaneous phase, with t ∈ [1, T]. The average PLV across the resulting matrix was used as a scalar estimate of the presynaptic synchronous activity, and it was incorporated into the MFM to modulate each K.

#### 2.1.3 Integration of axonal divergence

In SNN, axonal bifurcations were explicitly included in the microcircuit representation, especially when they present different synaptic conductance parameters, but this level of structural granularity was not preserved in the MFM derivation. Thus, in the MFM the contribution of axonal branches converging onto the same postsynaptic population was merged by a weighted summation of the corresponding synaptic parameters (see an example in section 2.3.1).

### 2.2 Construction of the Mean-field Model (MFM)

A second order MFM describes the activity of each population and the co-variances between population-activities with the following system of equations:

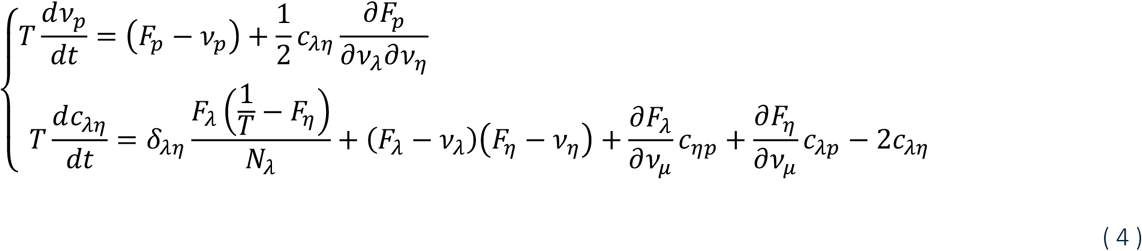

Where *v*_p_ = the activity of population p, c_λ*η*_ = the (co)variance between population λ and *η*, T = MFM time constant, N_λ_ = number of cells included in population λ, and F_p_ indicates the TF of population p.

Auto-MFM implements the MFM construction relying on the derivation of statistical moments describing the neuronal activity. The statistical moments are the average, the standard deviation and the autocorrelation time of membrane potential fluctuations and are used to parametrize the TF expression (F in equation 4), derived through an already defined and validated two step-fitting procedure^21^ that is summarized in the following section.

#### 2.2.1 Statistical moments of membrane potential fluctuations

Synaptic parameters are translated into the TFs via the presynaptic population specific conductance (*μ*_*Gs*_ [nS]) computed as:

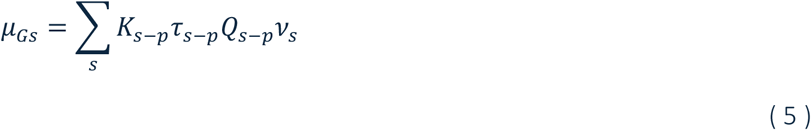

Where K_s-p_ = synaptic convergence weighted for the PLV, *τ*_s-p_ = synaptic decay times, Q_s-p_ = quantal conductance for each pair of presynaptic and postsynaptic populations (s-p), and *v*_*s*_ = presynaptic population activity ([Hz]).

The average population specific conductance was computed as:

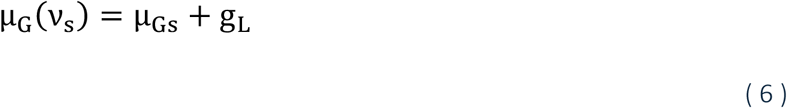

With g_L_ = leak conductance [nS].

The statistics of the membrane voltage potential are computed as function of µ_G_. In detail, the average membrane potential (µ_V_, [mV]) is:

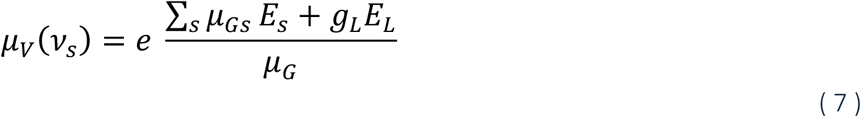

With E_s_ = reversal potential of the presynaptic connection [mV], E_L_ = rest potential of the postsynaptic population [mV], and µ_G_ = average conductance of target population [nS], while the variance (*σ*_*V*_ [*mV*]) and the autocorrelation time (*τ*_*V*_) of membrane potential fluctuations are expressed as:

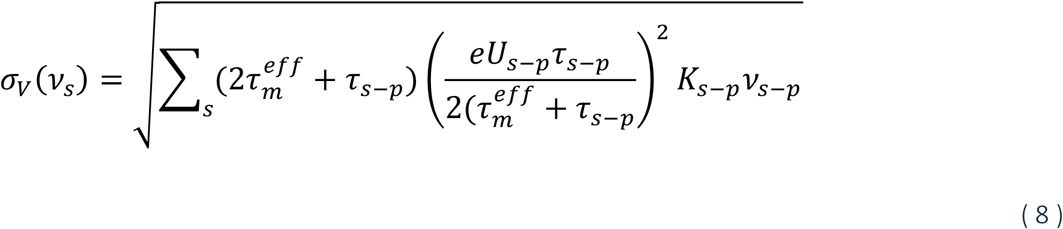

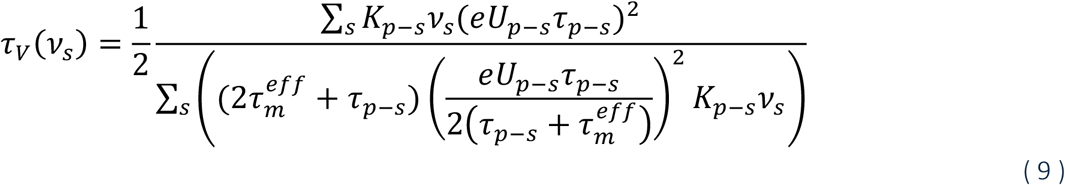

With 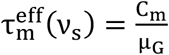, and 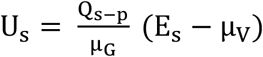

#### 2.2.2 Transfer function (TF) fitting

The derivation of the TF implemented in auto-MFM follows the semi-analytical approach detailed in^20^ and already validated for other MFMs, including the first version of the cerebellar MFM^25,31^.

In detail, the expression of a population-specific TF is:

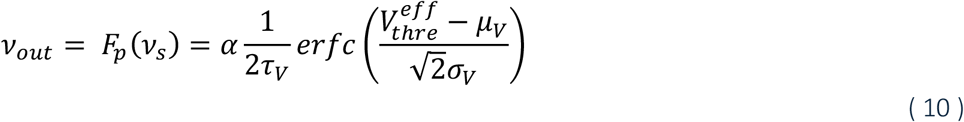

With ν_out_ = output activity, i.e., the firing rate modeled by the TF, F_s_ = TF of populations p in the equation, ν_s_ = presynaptic activity; α = phenomenological gain to capture both low and high frequencies, τ_V_ = autocorrelation time decay (ms), representing the timescale of the TF, µ_V_ = average membrane potential (mV), σ_V_ = standard deviation of the membrane potential (mV) and 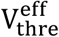= phenomenological threshold (mV), modeling the single neuron non-linearities. 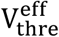 is expressed as a polynomial: in case of linear single-piont neuron model (e.g. LIF; E-GLIF), the expression is a first order polynomial, while for non-linear single-point neuron model (e.g. QIF; AdEx), it is a second order polynomial. Here it is reported the expression of 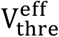 as a first order polynomial:

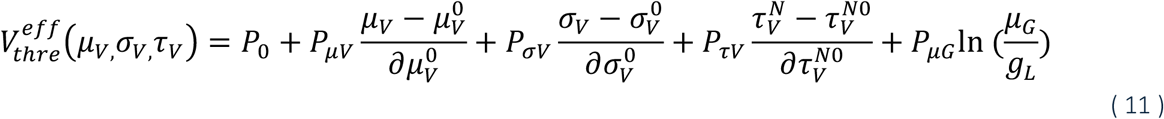

Where 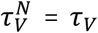 adjusted with the ratio between membrane capacitance and leak conductance 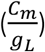, and 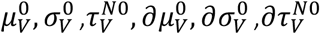 are rescaling constants to normalize the contribution of each term^21^. The coefficients P are computed using a double-step fitting yielding a numerical table (i.e., a numerical TF) containing the firing-frequency of the neural population extracted from the SNN^17^. The analytical TF is then obtained by fitting the P coefficients on the numerical TF.

#### 2.2.3 Optimization of the MFM

The phenomenological gain α of the TF (equation 10) enables MFM predictions both at low and high frequencies regime. For each population, α was optimized to match the input frequency-dependent response of the corresponding SNN neuronal population. Auto-MFM integrates a multi-objective optimization module based on the Non-dominated Sorting Genetic Algorithm (NSGA) (Python *pymoo* library), which is designed for multi-parameter problems^32^. In this framework, the population-specific α was treated as free parameter.

Two objective functions were defined:

1. Global error, to minimize the mismatch in magnitude between SNN and MFM activities. For each population and each input frequency, the error was computed as the difference between the average activity of the MFM and that of the SNN, normalized by the standard deviation of the SNN activity. Errors across populations were summed up to obtain an error score. Finally, these scores were aggregated across all the input frequencies to cover the entire physiological dynamic ranges.
2. Slope of the input-response relation of the output population, to give more relevance to its activity modulation.

The optimization was set as follows: 24 arrays, each one made up of a set of α values (one for each population) are generated at each iteration of the genetic algorithm, the maximum number of iterations is set at 50, tolerance = 10^-8^. The initialization was set equal to α values resulting from the fitting procedure. Higher penalization was assigned to the α combinations pushing the population activity out of its dynamical validity range. The optimization outcome is an ensemble of solutions, namely the non-dominated solutions, forming the Pareto front ^33^. The selection of the best parameters set amongst the non-dominated solutions was based on the knee-point detection, that is a method to select the solution corresponding to the maximum curvature of the Pareto front, i.e., the best trade-off of global error and slope of the input-response relation.

### 2.3 The cerebellar network test-bench

#### 2.3.1 Parameter-transfer from SNN to MFM

The main cerebellar neurons, i.e., GrC, GoC, stellate cells and basket cells, and PC, are wired in a multi-layer architecture: the cerebellar input stage is represented by GrCs and GoCs, both receiving excitatory input from mfs^34,35^. GrCs provide excitatory projections to GoCs, which in turn exert inhibitory feedback onto both GrCs and themselves, forming local recurrent loops. GrCs also project to the stellate and basket cells and PCs. Stellate and basket cells provide feedforward inhibition onto PCs, which constitute the sole output of the cerebellar cortex, projecting to the deep cerebellar nuclei. The biophysical-grounded SNN corresponds to a volume of the cerebellar cortex, of about 300 micrometers cube, including a total of ∼ 30.000 neurons. Each neuron type is represented by optimized EGLIF models (equation 1)^11,36^ whose parameters are tuned to reproduce in vivo microcircuit dynamics and are reported as supplementary material (Tab. S1 E-GLIF parameters and Tab. S2 cerebellar SNN parameters). The E-GLIF models are connected via conductance-based synapses (Eq. 2), with the conductance dynamics (g_syn_) set as an alpha function:

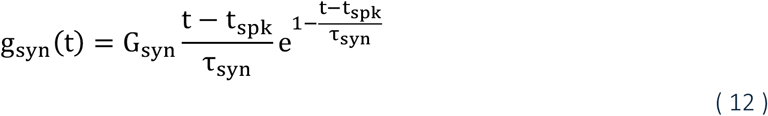

Where G_syn_ = conductance weight (nS), t_spk_ = time of a spike (ms), and τ_syn_ = synaptic time decay (ms), with all parameters specific for each connection type.

In detail, the biophysical cerebellar SNN was constructed using the Brain Scaffold Builder framework (BSB)^37^ which defines network topology using positioning rules and morphology-based connection rules, and interfaces with NEST to solve the dynamic equations^38^. From the SNN connectome, for each connection type, an average synaptic convergence (i.e., parameter K) was derived, and then it was scaled down according to the specific PLV of the presynaptic population, computed by imposing a 40-Hz input on mfs (representative of an intermediate frequency regime). Each connection type was functionally defined by setting a weight and a delay, representing the quantal synaptic conductance and the synaptic time decay (Q, *τ*).

The bifurcation of GrC axons into ascending axons and parallel fibers was explicitly represented in the SNN, with distinct synaptic parameters^39^. In the MFM, the functional parameters Q and *τ* of parallel fibers and ascending axons were averaged based on local synaptic density (i.e. number of synapses impinging on one representative PC), whereas the parameters K were scaled down according to the specific PLV of parallel fibers and ascending axons and subsequently summed.

As example for the connection GrC-PC, the excitatory synaptic parameters were computed as:

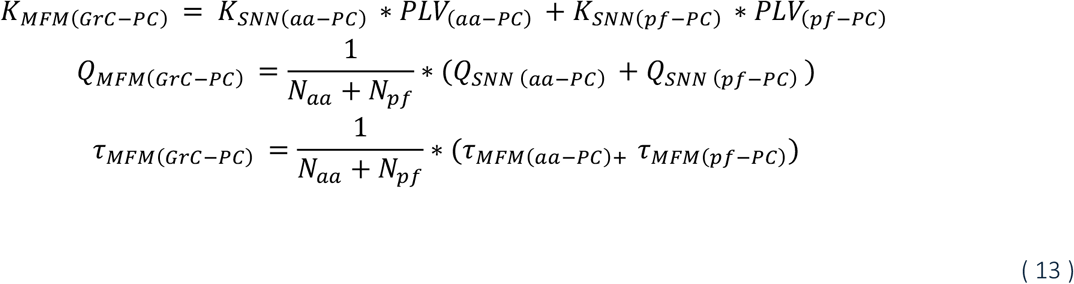

Where aa = ascending axons and pf = parallel fibers, and N is the number of incoming projections.

In more details, to quantify for instance the PLV (pf-PC), we extracted the firing rate signals from the 1500 parallel fibers converging onto a representative PC in the middle of the reconstructed volume and computed the pairwise PLVs across all presynaptic signals (see equation 3), resulting in a 1500 × 1500 matrix. The average of this matrix was used as a presynaptic population-level synchrony weight in the MFM connection parameters (K*PLV).

#### 2.3.2 Construction of the cerebellar MFM

The cerebellar MFM included the neuron types present in the cerebellar SNN as homogeneous populations. Stellate and basket cells were merged into a unique neuronal population, namely the molecular layer interneuron population (MLI) given their firing activity is similar, and they are involved in the same connection types. MLI synaptic parameters were derived by averaging the corresponding stellate and basket parameters and the transfer from micro to mesoscale was performed as in 2.1 consistently with all the other populations. The range of input frequency from mf bundles (*v*_drive_) spanned from 4 Hz to 80Hz, with a step of 4Hz and simulation lasted 5s (with a time step of 0.1 ms). PLV for all the synaptic connections was extracted for input frequency from mf (*v*_drive_) = 40 Hz and the population-specific TFs were computed^40^ (Fig.3). Numerical template for each population was extracted from the SNN as response frequencies. GoC population presents two excitatory inputs, one from mfs and one from GrC, and an inhibitory self-connection, therefore in addition to driving input from mfs, working frequencies were extracted for GrC and GoC itself. This yielded a 3D-TF that preserves the distinction between the two excitatory presynaptic sources, rather than collapsing them into a single aggregated excitatory drive.

**Figure 3.**
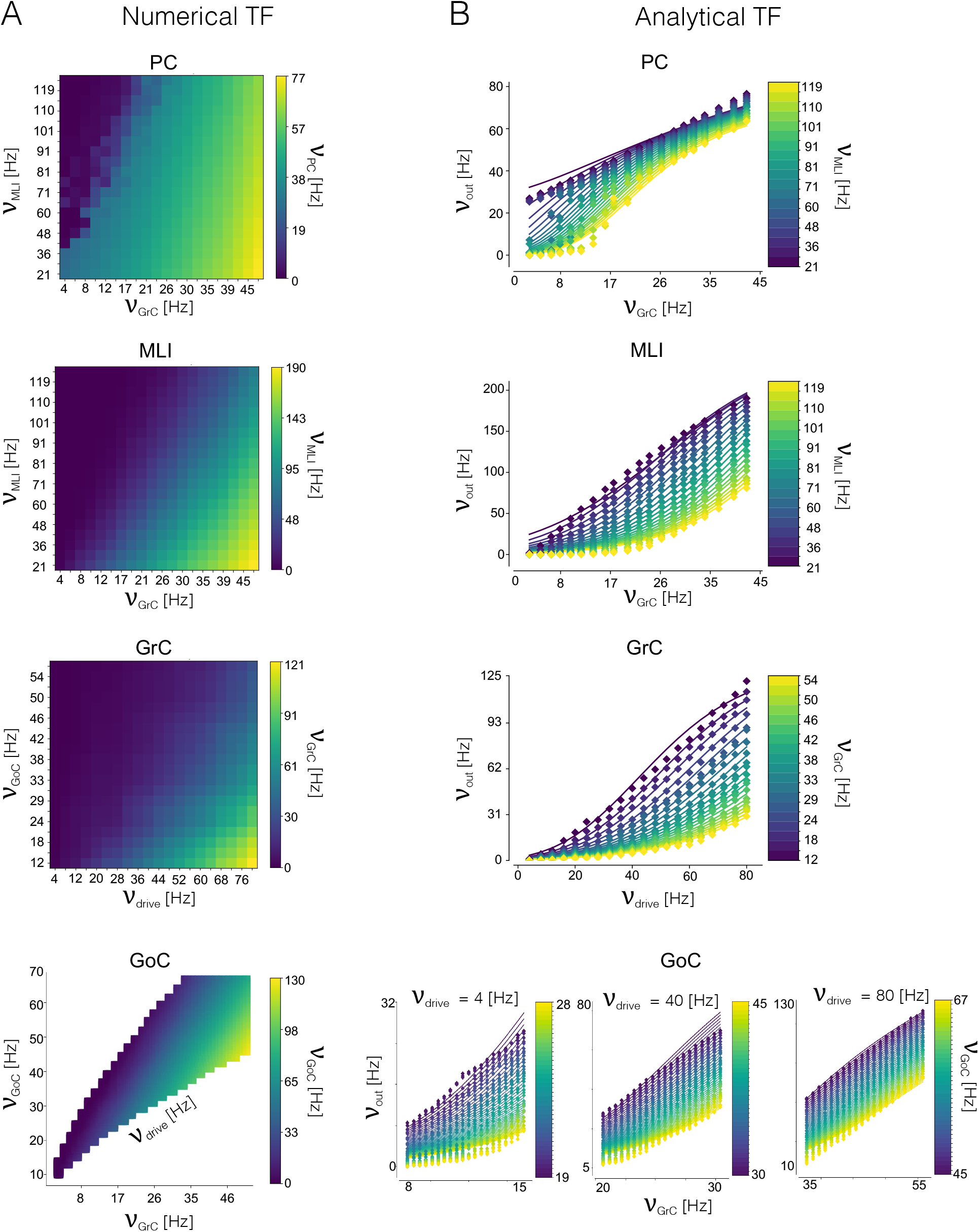
Transfer functions. Transfer functions (TFs) for the cerebellar MFM, tuned on the corresponding SNN, using the auto-MFM toolkit. **A)** Numerical TFs computed from SNN simulations for Purkinje Cells (PC), Molecular Layer Interneurons (MLI), Granule Cells (GrC), and Golgi Cells (GoC). Simulations lasted 5 s with a time step of 0.1 ms. Heatmaps report the output firing rate (color code) as a function of presynaptic input frequencies and, for GrCs and GoCs, also of input frequency imposed on mfs. **B)** Corresponding analytical TFs obtained by fitting the numerical data. Numerical TFs (points) are accurately captured by the curves (lines), demonstrating the robustness of the analytical TF fitting procedure across all neuronal populations (*v*_out_= firing rate estimated by the population-specific analytical TF, see Eq. 12). For GoCs, analytical 3D TF is shown at different mf drive frequencies (16,40,80 Hz), preserving the near-linear trend observed in the numerical domain.

Cerebellar MFM dynamical system integrates the population-specific TFs, as in equation 4. The cerebellar MFM time constant T is set at 3.5 ms, as estimated in Lorenzi et al., 2023^40^.

The optimization was performed on the four α of the TFs associated with GrC, GoC, MLI, and PC populations by minimizing the global error together with the slope constraint applied to PC population, i.e., the output of the cerebellar cortex. The optimization was run across noise-like input frequency of 4Hz, 40Hz, and 80 Hz. SNN and MFM activities were simulated for 500 ms, with a dt = 0.1 ms.

The optimization algorithm was run on a High-Performance Computing system using 32 core each and 256 Gb RAM and lasted 10 hours.

#### 2.3.3 Validation of Auto-MFM: cerebellar MFM-SNN responses

The auto-MFM tool underwent a constructive validation procedure to ensure that the MFM through parameters transfer, TF fitting and subsequent optimization preserved the firing activity of its microscale counterpart. For each neuronal population, the input-output relations of MFM and SNN were computed (simulation duration = 500 ms, dt = 0.1 ms) and compared in response to driving input from mf. For MFM the driving input was represented as a random-noise input. A set of 200 simulations were run for TFs construction to assess the robustness to stochasticity of the input. For each simulation, the mean ± standard deviation of MFM rate-based population activity was computed. For SNN, the driving input was defined as a Poisson spike train, and the activity was computed as #spike/seconds for each population.

Peristimulus time histograms (PSTHs; bin width: 10 ms; simulation duration = 500 ms) were computed for input frequencies of 40 Hz, representative of an intermediate frequency regime.

### 2.4 Exploitation of Auto-MFM to generate pathological variants

The flexibility of Auto-MFM in generating MFM was tested by deriving pathological variants of the cerebellar circuit. This was achieved by modifying specific synaptic parameters to emulate disease-related structural alterations (e.g., ataxia) or by constructing a model space spanning hypo-to-hyper excitability synaptic regimes relevant to different neurological and psychiatric conditions (e.g., hypoexcitation in schizophrenia, hyperexcitation in autism). Two possible scenarios for pathological MFM derivation were tested.

1. Pathology-driven modification. When a pathology is known to alter specific circuit parameters, Auto-MFM was used to automatically generate the corresponding pathological MFM variant by incorporating the altered parameter value into the construction pipeline.
2. Model parameter space. When the focus is on the effects of a pathophysiological variation of a parameter, Auto-MFM was used to generate a model library by sweeping the relevant parameter range.

#### 2.4.1 Purkinje Cells (PC) dendritic simplification (ataxia-driven modification)

Ataxia is associated with a reduction of the PC dendritic tree, which leads to a decreased synaptic density. In SNN, the reduced complexity of dendritic branches directly impacts in the number of synapses found by the morphology-based connectivity rules. The parameters *K* extracted from the SNN reconstruction are reported in Table 1, and the level of PC dendritic simplification was set to match experimentally observed reduction of GrC-PC synapses of ∼ 25% (ref). These *K* values, related to synaptic convergence on PCs, were then passed in the MFM parametrization resulting in:

**Table 1.** PLV between presynaptic and postsynaptic populations in the cerebellar SNN. PLV quantifies the degree of phase synchronization between presynaptic spike trains converging on a given postsynaptic target and is used as a weighting factor to reparametrize the synaptic convergence (i.e., *K*_*s-p*_) when scaling from the microscopic to the mesoscopic domain. mf = mossy fibers, pf = parallel fibers, aa = ascending axons.

| Presynaptic | Postsynaptic | PLV |
| --- | --- | --- |
| mf | GrC | 0.506 |
| mf | GoC | 0.350 |
| pf | GoC | 0.358 |
| pf | Basket | 0.398 |
| pf | Stellate | 0.358 |
| pf | PC | 0.358 |
| aa | GoC | 0.633 |
| aa | PC | 0.460 |
| GoC | GoC | 0.453 |
| Basket | Basket | 0.402 |
| Basket | PC | 0.419 |
| Stellate | Stellate | 0.413 |
| Stellate | PC | 0.661 |

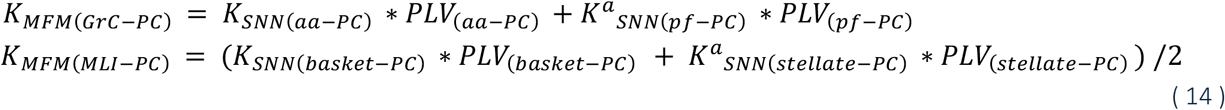

Where K^a^_SNN_ are the reduced synaptic convergences resulting from ataxia-like configuration: *K*^*a*^_*SNN(pf−PC)*_ = 1110 (i.e., reduction of 26% compared to control values); *K*^*a*^_*SNN(stellate−PC)*_ = 3.7 (i.e., reduction of 21% compared to control configuration). K_SNN_ are set at control configuration values: *K*_*SNN(aa−PC)*_ = 92, *K*_*SNN(basket−PC)*_ = 19. MFM synaptic parameters are: *K*_*MFM(GrC−PC)*_ = 439.7 and *K*_*MFM(MLI−PC)*_ =3.75 Dendritic simplification does not affect aa-PC synapses for geometrical constraint, nor basket-PC since the synapses are between the soma rather than the dendritic tree. Auto-MFM was used to derive the PC TF using these parameters K, while Q and *τ* are set as in control configuration (section 2.2). The TF parameter *α*_*PC*_ was further tuned to match a reduction of PC baseline activity in the case of spinocerebellar ataxias^41–43^. Specifically, *α*_*PC*_ was tuned on ∼ 30% reduced PC firing in basal condition^42^, corresponding to a reduction in simulated PC activity from ∼ 95 Hz to ∼ 65 Hz in basal condition (i.e., input from mfs = 4Hz). An input space exploration was performed by applying a transient external perturbation to the MFM in the form of a Gaussian impulse from mossy fibres (*σ*= 0.01). The impulse amplitude was systematically varied within the range [4:4:80] Hz, and the resulting peak of PC dynamics were extracted. Statistical differences between control and ataxia-like conditions were evaluated using the Wilcoxon signed-rank test with significance level. Additionally, slope differences between control and ataxia-like linear fits were computed to assess alterations in PC signal propagation. Paired comparisons across input levels were performed on PC simulated activities using a Wilcoxon signed-rank test (significance threshold *α* = 0.05). The relationship between input amplitude and peak response was approximated using linear regression, yielding a best-fit line for each population. The quality of the linear approximation was quantified using the coefficient of determination (R^2^) and the slope differences between control and ataxia-like linear fits were computed to assess alterations in PC peak.

#### 2.4.2 Model-based space of hypo-to-hyper excitation of granular layer (parameter-driven exploration)

A systematic parameter sweep was performed on the mf-GrC transmission (i.e., excitatory quantal conductance parameter Q), spanning different physiopathological states, starting from the standard value Q_mf_GrC_ = 0.23 nS and exploring a range from 0 to 2× Q_mf_GrC_, with a step of 10% (Tab. S3 for the parameter sweep). All the other synaptic parameters were kept at their standard configuration. The auto-MFM pipeline was executed to generate a taxonomy of GrC TFs, and consequently a taxonomy of MFMs, so defining a model-based space. Reduction of Q_mf_GrC_ might represent a proxy of impaired transmission of excitatory signals form cerebral cortex (via mfs) to the cerebellar cortex, such as in cerebellar-related schizophrenia, while increasing of Q_mf_GrC_ reproduced GrC hyper-excitability observed in pathology as autism. Experimental recordings in a mouse model of autism showed a 60% of increase in NMDA conductance in GrCs^44^. To obtain the equivalent MFM, the GrC TF corresponding to Q_mf_GrC_ +60% (0.37 nS) was selected from the taxonomy. The resulting dynamics were analysed across all network layers. For each population and input frequency, the peak values of the MFM activities were extracted. The same analysis used for ataxia-like case was applied to quantify the differences of signals propagation between control and autism-like conditions.

## 3. Results

Auto-MFM proved an efficient framework to automate the generation of a MFM from a SNN. Auto-MFM was tested on the cerebellar SNN, achieving a flexible and automatic corresponding MFM^28,45^ validated against single-neuron-resolution spiking signals. Furthermore, pathological variants were defined from structural or functional neural alterations, focusing on one pathology-related value, or exploring the consequences at network level of one MFM parameter space.

### 3.1 Cerebellar MFM construction and validation

PLV scores (Tab.1) indicated that effective synaptic transmission synchrony reduced structural synaptic convergence (K) values by a factor ranging from 0.35 to 0.66. The result of the entire parameter-transfer pipeline (section 2.1) is reported in Fig 4. In the MFM implementation, stellate and basket cells collapsed into MLI as a unified molecular layer interneurons population, with their PLV values averaged and weighted by the respective number of cells in the generative SNN.

**Figure 4.**
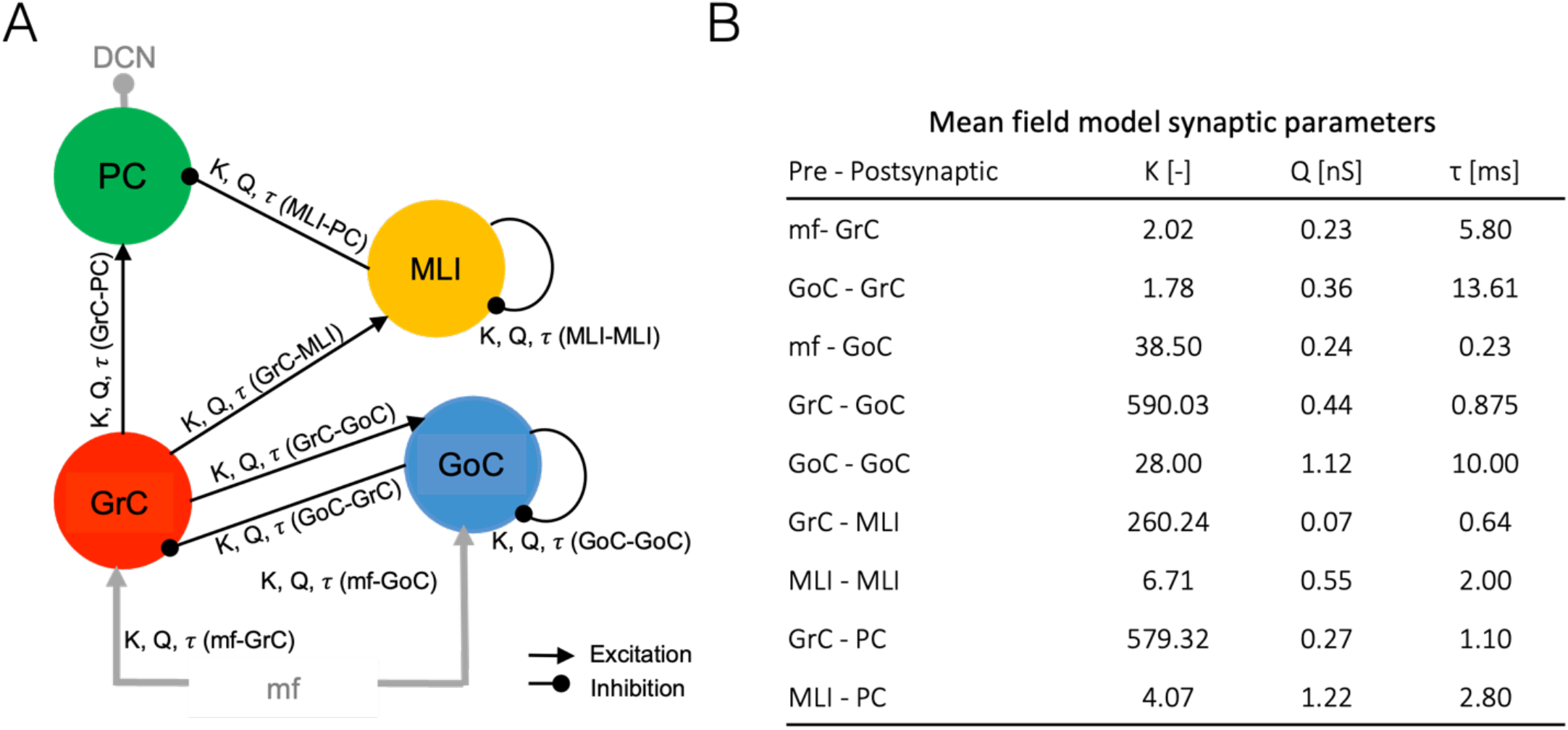
Cerebellar MFM configuration. **A)** Multi-layer network of the cerebellar MFM, with the synaptic parameters specific for each connection type, extracted from biophysical-grounded SNN and tuned for population-level formalism, using Auto-MFM. **B)** Cerebellar MFM synaptic parameters. The synaptic convergence K (number of presynaptic neurons per postsynaptic neuron) was calculated from the anatomical convergence in the SNN multiplied for the synaptic-specific PLV value (see Tab 1), the quantal synaptic conductance Q was derived from the synaptic weights of the corresponding SNN synapse models, while the synaptic time constant τ was taken from the decay constants of the SNN synaptic dynamics. Re-parametrization was computed as in Eq. 6. K values reported are multiplied by the corresponding PLV values. As for the SNN, the K values for autoinhibition (i.e., K_mli-mli_ and K_goc-goc_) were tuned to ensure physiologically plausible baseline firing rates in the MFM (De Schepper et al., 2022).

**Figure 4.**
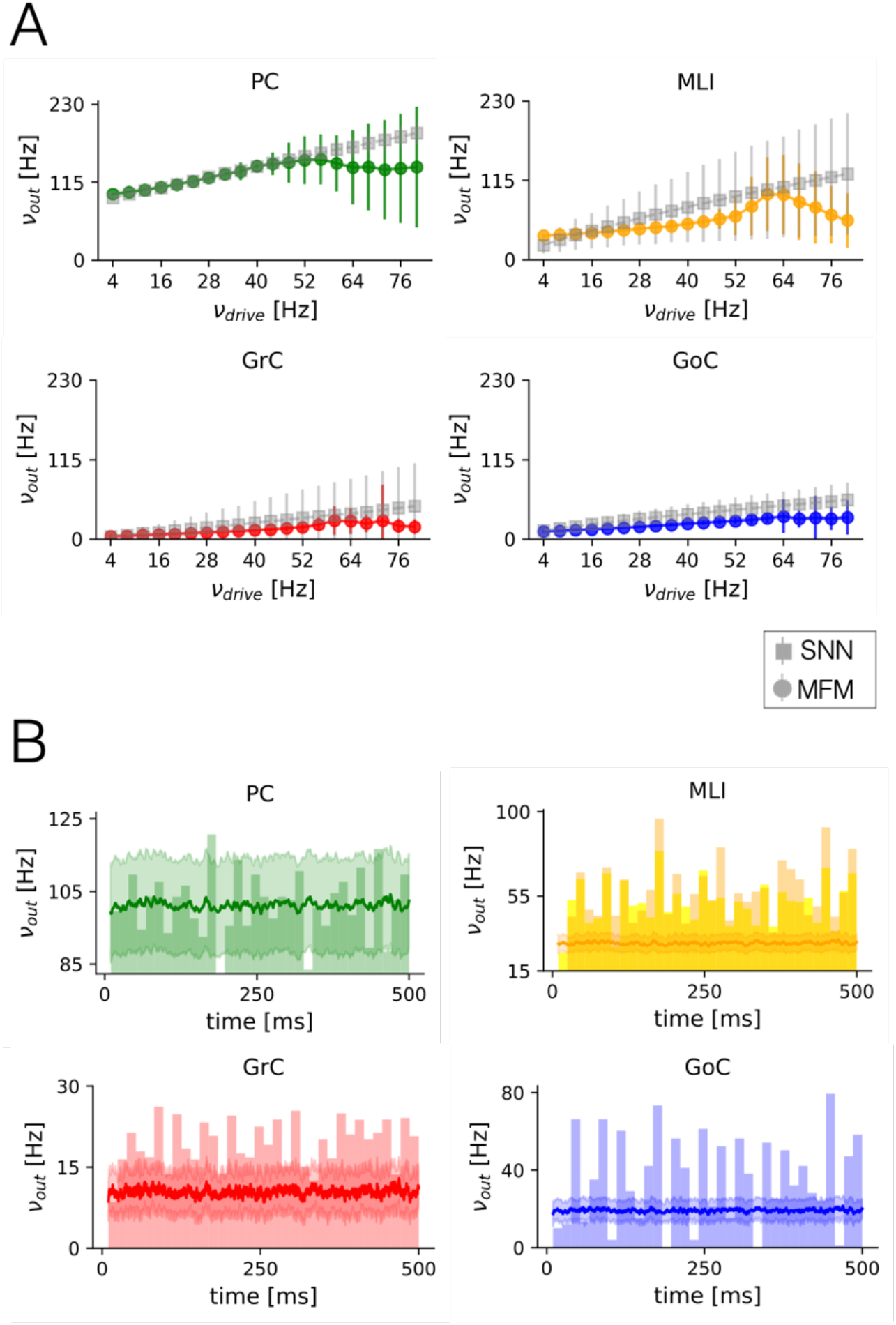
Comparison of MFM and SNN. **A)** Input/output relation of MFM. 100 simulations of random noise-like input were performed for each input frequency used in the construction of Numerical TF (range = 4:4:80 Hz). The resulting MFM activity was averaged across simulations (colored dots = mean activity; colored lines = standard deviation) and overlapped to SNN simulations under the same conditions (gray squared = mean activity; gray lines = standard deviation). MFM activity results in the standard deviation of SNN for all the population, with PC reproducing the SNN output almost exactly for inputs between 4 and 60 Hz. At higher input frequencies, the stochasticity in the definition of the noise introduces fluctuations in the averaged MFM-predicted activity. These fluctuations remain within the standard deviation of the SNN output, but the resulting trend deviates from SNN linear input/output relation. **B)** Cerebellar MFM activity for a random noise-like input (*v*drive, from mossy fibers) at 40 Hz overlayed to the corresponding PSTH extracted from SNN (PC = green, MLI = orange, GoC = blue, GrC = red; MFM activity = solid line, standard deviation = shadow; SNN PSTH bin = 10 ms). For the neuronal populations the MFM rate-based signal represents the average of PSTHs, with best matching for PC. Only for MLI the activity simulated with MFM is reduced compared to the stellate and basket cells PSTHs

The multi-objective optimization showed that, in the final Pareto front, the α parameter set presented a monotonic-decreasing distribution, suggesting a reliable performance of the algorithm (Fig. S1). 13 out of 24 arrays survived as non-dominated solutions (i.e., 13 combinations of [α_GrC_, α_GoC_, α_MLI_, α_PC]_). For all of them, the global error was more sensitive to different α parameter sets included in the Pareto front than the output slope. This result is due, first, to the fact that the global error is an aggregate metric computed directly over the activity of each population, while the slope (of PC activity) directly influences only α_PC._ Second, the slope reflects a trend rather than a magnitude, and the starting point was already close to a linear input–output relationship. In this regime, the score provides finer granularity, as even small slope variations are detected more easily than comparable changes in activity magnitude. The α values initially set for TF fitting were α_GrC_ = 2.6, α_GoC_ = 1.9, α_MLI_ = 1.8, and α_PC_ = 1.8. The best solution identified by the knee algorithm resulted in α_GrC_ = 2.1, α_GoC_ = 2.4, α_MLI_ = 1.6, and α_PC_ = 5.4, showing that PC TF required a substantially higher phenomenological gain compared to the other cerebellar populations.

The MFM constructed with the parameter-transfer and optimization procedures included in Auto-MFM showed activities that matched the generative SNN ones (Fig. 5). Indeed, activity simulated with the MFM fell in the standard deviation of the one simulated with the SNN (Fig. 5A). For GrC, GoC and PC populations, the input/output relation of the MFM matched exactly the relation of the SNN till input frequency of about 60 Hz, retracing the linear trend of SNN activity. For higher noisy input on mfs, small differences in GrC population emerged, and reverberated mainly in MLI trend. The PSTHs showing the distribution of the spiking activity for mfs input = 40 Hz are overlaid with the rate-based signals produced by the MFM under the same input pattern (Fig. S2), providing a qualitative comparison between the different domain of SNN and MFM (Fig. 5B). Although some differences due to single-spiking-neuron-resolution signals vs rate-based averages, PC activity closely matched the SNN response, also thanks to a dedicated objective in the optimization procedure. This population-specific slope optimization was not included for all populations to prevent a growing computational load in contrast with the intended use of the MFM approximation. Overall, MFM preserves the effective input-output relation of the whole multi-layer circuit with an internal signal transmission slightly different from the microscale SNN, due to the distinct operational domain of SNN and MFM that required a parameters manipulation to move from one to another domain (e.g., synchrony weighting for PLV, or merging of axonal bifurcations).

**Figure 5.**
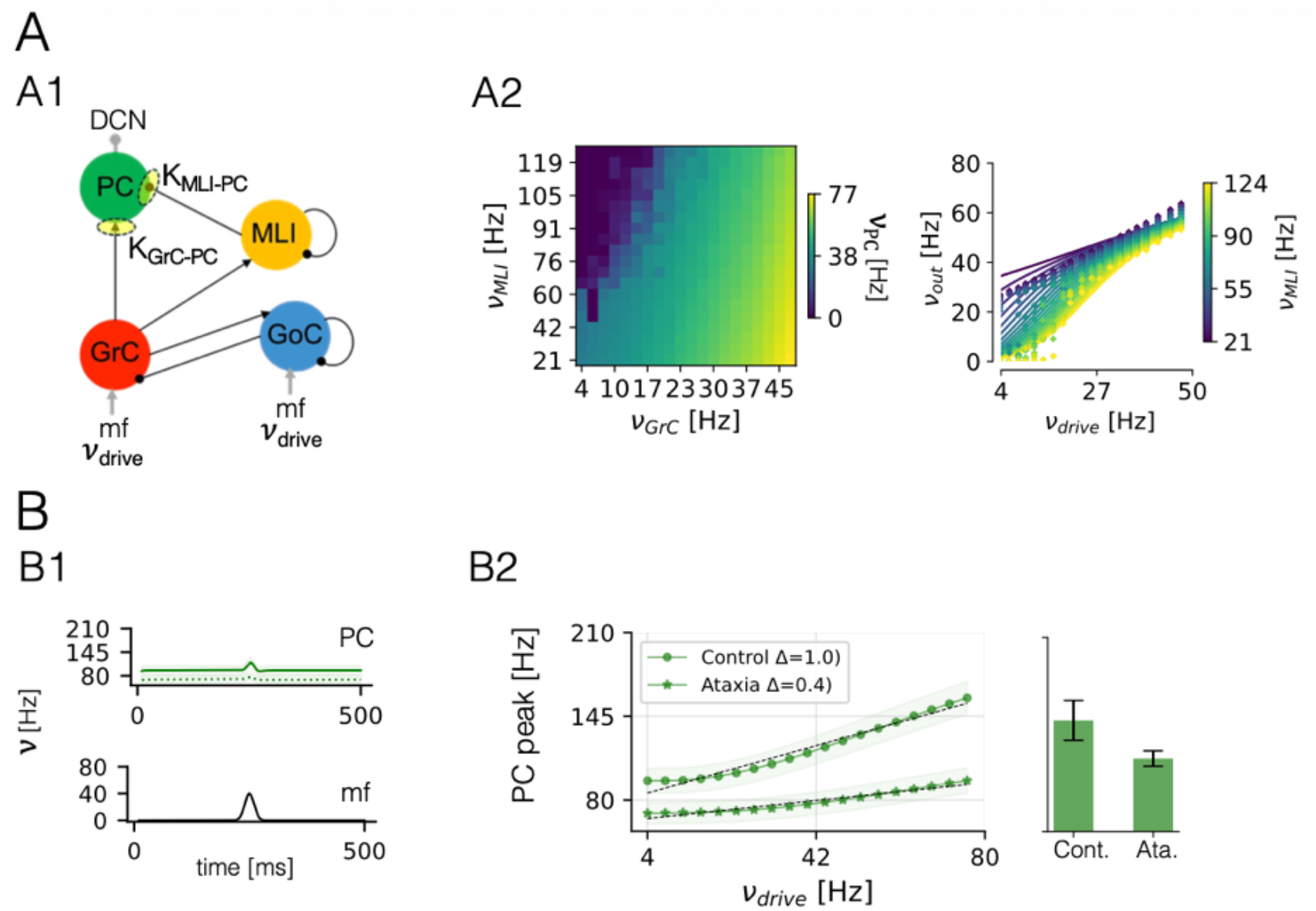
Ataxia-like circuit modification. **A)** Ataxic MFM derivation. **A1)** Effect of ataxia modeled as PC dendritic simplification, parametrized through the reduction of excitatory and inhibitory synaptic convergence to PC (i.e., K_GrC-PC_ and K_MLI-PC_, respectively). **A2)** Auto-MFM tool is used to derive the ataxic PC TF (numerical template and the correspondent fitting of the analytical TF) which shows a reduced firing rate (*v* ™ [Hz]) compared to the firing rate of control PC. **B)** Ataxic MFM prediction. **B1)** PC activity in control case (continuous line) and ataxia configuration (dotted line) in response to a fast-dynamics stimulation from mf (*v*_drive_: gaussian shape, amplitude = 40Hz, sigma = 0.01). **B2)** PC peak computed across input frequency values. PC peak shows a quasi-linear trend (control PC = dots, ataxic PC = stars, standard deviation as shadow). PC peak is significantly reduced in ataxic PC compared to control one (Wilcoxon-test, significance threshold = 0.05, p-value < 0.001). The slope of *v*drive-PC peak relation is steeper in control PC, showing that ataxia slows down both the amplitude of the PC peak and the sensitivity to input intensity. PC peak averaged over the input frequency space (bar plot), shows the overall difference of peak amplitude between control (Cont.) and ataxic (Ata.) configurations.

### 3.2 Cerebellar pathology-like variants

#### 3.2.1 Purkinje Cells (PC) dendritic simplification: ataxia-like circuit modification

The analytical PC TF in the ataxia-like condition (Fig. 6A) preserves the overall shape observed in the control case (see Fig. 3), but its amplitude is reduced. This attenuation reflects the stronger decrease of the excitatory convergence *K*_*GrC–PC*_ (∼ 24% reduction, considering the aa and pf synapses) compared to the milder reduction of the inhibitory convergence *K*_*MLI–PC*_ (∼ 8% reduction). As a consequence, excitation remains predominant over inhibition (i.e., monotonic increasing relation between PC firing rate and GrC input), but with lower magnitude. In the MFM predictions, ataxic PC activity is significantly reduced relative to control PCs (Wilcoxon test, p < 0.05), with the largest differences emerging at higher input frequencies. Control PCs exhibit a steeper increase in firing rate as input frequency rises. This divergence is quantified by the slope of the PC input–output relation with increasing mf inputs. Control PC shows a slope (Δ) of 0.23, whereas the ataxic PC displays a reduced Δ of 0.08. Therefore, ataxic PCs are not only reduced in firing magnitude but also compromised in transmission fidelity, since a poor modulation of their response based on input intensity

**Figure 6.**
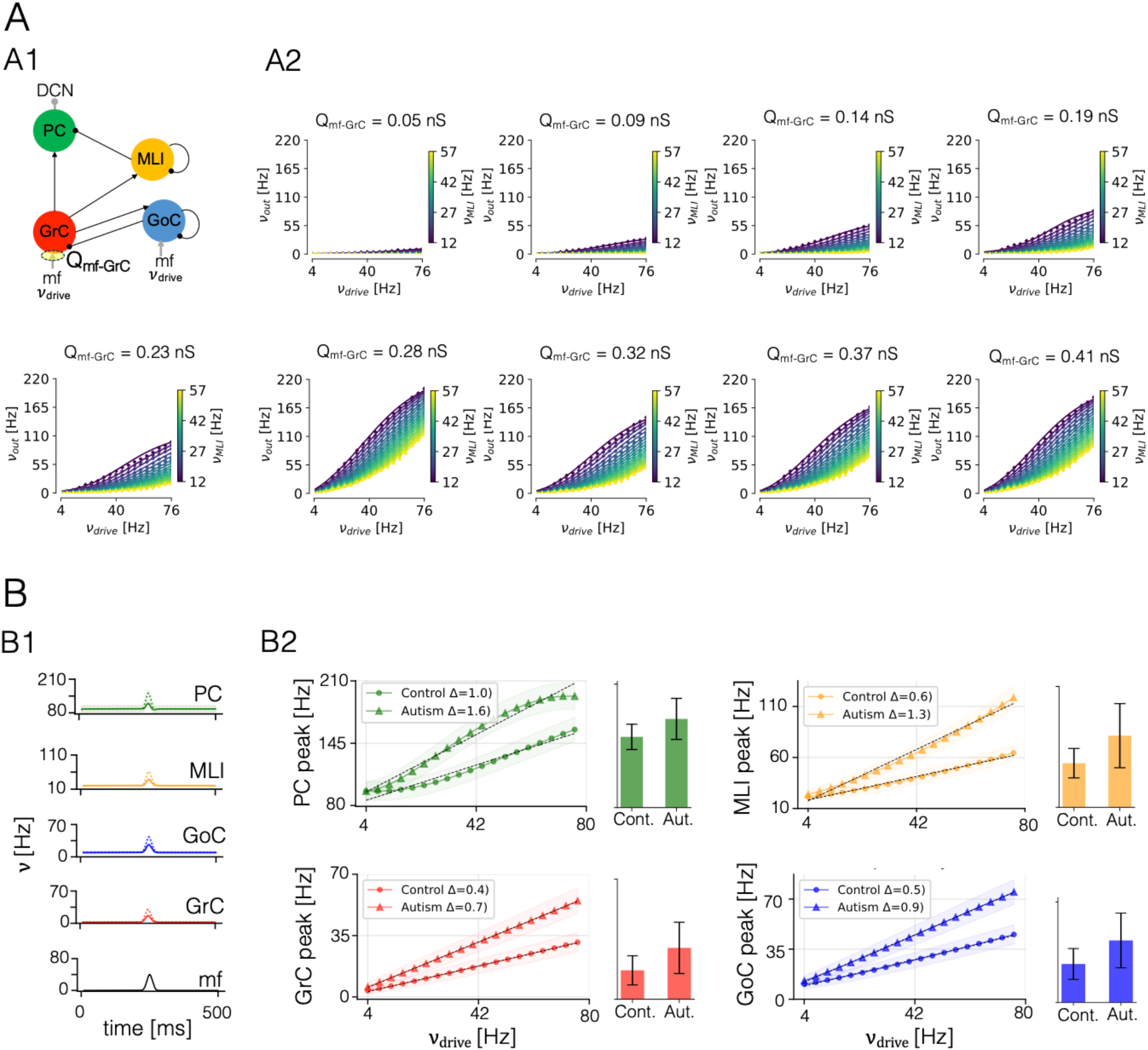
Exploration of model space through a variation of the excitatory synaptic conductance. **A)** GrC TF taxonomy. **A1)** Excitatory synaptic conductance from mf to GrC (Q_mf-GrC_) is systematically varied respect to its baseline value (= 0.23 nS) resulting in hypo and hyperexcitable configuration. **A2)** Analytical TFs are computed following the two-step fitting procedure for all Q_mf-GrC_ values. Here only the variation of 20% are reported, complete taxonomy in supplementary material (*v*_drive_ = excitatory input from mfs [Hz], *v*_GoC_ = inhibitory input from GoC [Hz], *v*_out_= GrC activity [z]). GrC analytical TFs capture the fast-rise dynamics for increasing Q_mf-GrC_, and a progressively reduction of the GoC inhibitory effects onto GrC dynamics. **B)** MFM prediction in autism (Q_mf-GrC_ = 0.37 nS, increase of 60%) **B1)** MFM activity in response to a fast-dynamics stimulation from mf (*v*_drive_: gaussian shape, amplitude = 40Hz, sigma = 0.01). Control configuration (continuous line) and autism-like configuration (dotted line). Hyperexcitability of GrC is propagated over the forward layer. **B2)** Population peaks computed across input frequency space (control = dots, autism = triangles, standard deviation as shadow). The peak activity is significantly increased for all the population in autism configuration compared to control one (Wilcoxon-test, significance threshold = 0.05, p value < 0.001). The slope of PC peak-*v*drive relation changes the turned over the input frequency, with a tendence to reach a plateau for higher input frequency. Population peaks averaged over the input frequency space (bar plot), shows the overall difference of peak amplitude between control (Cont.) and autism (Aut.) configuration, with a reduced difference in PC cells, comparing to the other population, due to the difference in of PC peak-*v*drive relation trend.

#### 3.2.2 Model-based space of hypo-to-hyper excitation of granular layer

Auto-MFM was used to define a model space of GrC activity, that is transmitted to the forward layers (Fig. 7). The taxonomy of GrC TFs revealed distinct dynamical regimes captured by the GrC TF fitting (Fig. S3 and S4). GrC analytical TFs (Fig. 7A - continuous lines) captured the trend of the numerical TF (Fig. 7A – dots). GrC analytical TFs showed a high dependence on the magnitude of the excitatory conductance alteration. GrC TF computed in standard configuration (Q_mf-GrC_ = 0.23 nS), exhibits a gradual increase in output firing rate as mf input frequency rises, with the response within 125 Hz for lowest inhibition (from GoC) and highest excitation (from mf). When excitatory conductance was reduced below the standard configuration, the GrC TF compressed (e.g., *v*_out_ was lower than 120 Hz, for lowest inhibition and highest excitation, for all the reduced value Q_mf-GrC_ tested). This attenuation impaired high-frequency responsiveness, reproducing a hypoactive regime that could compromise signal propagation to the molecular layer. Under hyperexcitation condition, the GrC TF presented a strong amplification (e.g., Q_mf-GrC_ +60%, *v*_out_ is up to 240 Hz, for lowest inhibition and highest excitation), producing steeper input–output curves that model the GrC hypersensitivity to the excitation projecting from mfs. MFM activities under different input patterns were simulated for with Q_mf-GrC_ +60% to reproduce GrC hyperexcitation observed in autism^44,46^ . This analysis enabled assessment of whether hyperexcitation effects could be described as a uniform gain modulation across populations or whether layer-specific deviations were present. Using a fast-pulse stimulation protocol, the maximal slope of the input– output relation for each population was quantified in the autism-like configuration (Q_mf–GrC_+60% = 0.37 nS). All populations exhibited a steeper increase compared to control: granule cells (0.4 vs. 0.7; difference = 0.3), Golgi cells (0.5 vs. 0.9; difference = 0.4), molecular layer interneurons (0.6 vs. 1.46; difference = 0.9), and Purkinje cells (1.0 vs. 1.6; difference = 0.6). These results indicate a faster escalation of firing-rate responses across the circuit under the autism-like condition, with a higher impact in molecular layer. Moreover, the relation between PC peak and *v*_*drive*_ show a difference both in magnitude (increased in autism configuration) but also in the trend. Indeed, while all the other populations maintain a linear trend, PC show the tendence to reach a plateau for higher input frequency. A difference in the excitatory/inhibitory balance of PC is suggested with hyperexcitability of GrC triggering an over inhibition from MLI to PC, that modify the PC response in input frequency space.

## 4. Discussion

The auto-MFM tool developed here is a novel computational framework for automated multiscale modeling of brain circuits, designed to streamline the generation of mesoscale models (i.e., MFMs) from their biophysically-based microcircuit counterparts (i.e., SNNs). In essence, Auto-MFM leverages on two fundamental assets: 1) the MFMs construction from biophysical SNNs, achieved thanks to a fully automated translation of parameters from micro-to-mesoscopic domain; 2) the transformation of a MFM into pathological variants.

Auto-MFM demonstrated to facilitate the construction of a complex multi-layer MFM such as the cerebellar one, while minimizing manual intervention during parameter-transfer and preserving microcircuit features often lost in population-level simplifications. In the SNN, indeed, the connectivity of each point-neuron is probabilistically defined based on its 3D spatial location and inward and outward projection rules, thus providing heterogeneous transmission pathways. Since the MFM formalism collapses neurons into homogeneous populations, that microstructural heterogeneity is lost and an automated parameter-transfer strategy needs to be carefully defined. For instance, Auto-MFM estimates the value of synaptic convergence by weighting the structural convergence with an average synchronization of signals (i.e., quantified as the PLV) from axons projecting to common postsynaptic targets. Moreover, replacing the previous manual trial-and-error tuning of TF gain parameters (i.e., *α*) with a multi-objective genetic algorithm allows to increase reproducibility and robustness.

### 4.1. Validation of Auto-MFM: MFM-SNN responses

While procedural advantages are clear in facilitating any MFM construction from corresponding generative SNNs, Auto-MFM was specifically tested to prove physiologically reliability of a complex MFM of the cerebellar cortex, retaining both multi-layer architecture and population-specificity of TFs. Indeed, the automatization of MFM derivation through a seamless parameter-transfer from SNN to MFM and optimization of TFs is crucial to speed up the generation on new mesoscale representations of brain area, but the physiological reliability of this procedure must be ensured to provide an effective neuroscientific tool.

Given its structural and functional complexity, the cerebellar MFM represents a challenging test-bench for the framework. Auto-MFM has been used to update the cerebellar MFM previously published^40^ by incorporating synchrony-driven parameter-transfer and automated optimization of the population-specific TFs. The underlying SNN represented the mouse cerebellar canonical circuit and was refined to specifically reflect the physiological in-vivo states, improving the physiological relevance of MFM in the perspective to be used in whole-brain simulators, which in-silico reproduce in-vivo brain dynamics^47–49^.

It is worth noticing that the MFM is not intended to replace the SNN. They are complementary approaches to study the complexity of neuronal dynamics in a multiscale pipeline: SNN captures the precise timing and interactions of multiple single neurons, while MFM offers a higher-level representation of average firing rates and population dynamics. The predicted signals thus lay in different domains: SNN reproduces the activity in terms of number of spikes in a time window, while the MFM provides rate-based signals continuously modulated in time. Nonetheless, Auto-MFM pipeline preserves salient dynamics features. The cerebellar MFM faithfully rate-codes the corresponding SNN activity over a broad range of input frequencies (Fig. 5). At the cerebellar network output stage, PCs show an almost identical activity pattern in the MFM as in the SNN over the 4–60 Hz input range, while remaining within the standard deviation of the SNN response at higher-frequency inputs. It should be noted that the MFM output (i.e., the PC for the cerebellar cortex) undergoes a more intense TF optimization than other populations, with a dedicated objective of the optimization algorithm (i.e., the slope of the input/output relation) because its physiological reliability is crucial when a MFM becomes a functional unit in a virtual brain. When integrated into whole-brain simulators, the cerebellar cortex output generated by PCs is connected to the Deep Cerebellar Nuclei (DCNs), that, in turns, project to the thalamus and the cerebral cortex. The inhibitory synaptic connectivity between PC and DCN plays a crucial role in complex mechanisms such as long-term depression/potentiation^47^ and impacts on whole-brain dynamics^47,50^. In aggregate, an automated framework capable of translating microscale circuit features into high-fidelity mesoscale models provides a principled mechanism to systematically convert spiking neural networks into mean-field representations. In this perspective, Auto-MFM enables individual microcircuits to be seamlessly integrated as functional building blocks of a virtual brain, effectively bridging microscale mechanisms and macroscale brain dynamics through accurate large-scale simulations.

### 4.2 Auto-MFM to simulate circuit variants and pathological changes

The full potential of accurate large-scale simulations emerges when simulating pathological conditions. Given the heterogeneity of pathological mechanisms, having a reproducible and reliable computational workflow maintaining salient architectural and electrophysiological features is critical. Auto-MFM can directly support physiological validation and the controlled exploration of circuit variants in different pathophysiological conditions, linking local changes (e.g., mfs-to-GrC conductance) to emergent population responses in terms of firing rate activity. Therefore, Auto-MFM, by enabling the derivation of pathological MFMs, facilitates also the construction of virtual brains informed by specific neuronal and synaptic alterations. Virtual brains, that use MFMs as region models^47,48,51^, can thus be used to assess the impact of neuronal population specific synaptic alterations onto large-scale signals, like those recorded using functional magnetic resonance imaging or electroencephalography.

#### 4.2.1 Purkinje Cells (PC) dendritic simplification: ataxia-like circuit modification

Experimental recordings in mouse models of ataxia show a reduction in dendritic spine development in PCs^49^. The ataxia-like modification implemented in our model, by differently reducing *K*_GrC-PC_ and *K*_MLI-PC_, mainly captures the decreased PC ability to integrate excitatory inputs from granule cells. Consistent with this, experimental studies report decreases in parallel-fiber synaptic markers such as VGLUT1, alterations in postsynaptic glutamatergic signaling, and reductions in dendritic spine density or dendritic arbor length across multiple ataxia models^55,56^. Although the magnitude of these alterations varies across etiologies, their functional impact converges toward a diminished excitatory drive and a restricted dynamic range of PC responses. The cerebellar MFM reduction reproduces this principle: the attenuated slopes in the *v*_drive_– *v*_PC_ relationships reflect a reduced response modulation to changes in input frequency. This pattern is compatible with a limited capacity of PCs to acquire and consolidate afferent excitatory inputs and with the emergence of a more rigid, less adaptive movement-related output, a hallmark of the motor impairments typical of cerebellar ataxias.

#### 4.2.2 Model-based space of hypo-to-hyper excitation of the granular layer

Auto-MFM enables the simulation of circuital changes localized at a specific synaptic connection through a seamless derivation of a population-specific TF taxonomy, representing the model space where different pathological variants live. The model space identified here for a sweep of Q_mf-GrC_ parameter covers the pathophysiological conditions ranging from hypo-to-hyper excitability, as reported in schizophrenia and autism ^44,57–59^. The TF taxonomy can be exploited in two complementary ways. In the forward direction, when a pathology is associated to a certain parameter value, the corresponding TF can be selected directly from the taxonomy to generate a complete MFM that incorporates that specific modification. In the inverse direction, when the alteration is observed at the level of population firing rate, the taxonomy enables the identification of the underlying parameter change that best reproduces the recorded activity. In both cases, selecting a pre-fitted TF allows the MFM to emulate how a given condition of mf–to-GrC synaptic transmission impacts at cerebellar network level.

An example of the forward use of the taxonomy is provided by the autistic IB2-KO mouse model, in which an increased NMDA conductance has been experimentally reported ^44^. The hyper-function of NMDA in granular layer is estimated in an 60% increase of Q_mf–GrC_ . The corresponding TF predicts an overall enhancement of activity across all other populations, indicating how an excitatory–inhibitory imbalance at the input stage propagates through the circuit.

An example of the inverse use exploits TFs associated with GrC hypo-excitability, a condition relevant to several neuropathologies. For instance, schizophrenia is characterized by glutamatergic synaptophysin, where NMDA receptor hypofunction and associated synaptic deficits disrupt the excitation–inhibition balance^57^. Firing data from granule cells in schizophrenia would drive the selection of the “schizophrenic” granular TF, which then would be inserted in the cerebellar MFM. Although a MFM for this specific case was not derived here, the TF taxonomy indicates that reduced GrC excitability would propagate forward as a global decrease in activity across subsequent populations, consistent with impaired information flow through the granular layer.

In both cases, with the cerebellar MFM, it is possible to simulate the propagation of GrC activity over the layers to generate knowledge about the excitatory/inhibitory balance of the whole network. Overall, Auto-MFM opens the possibility for a seamless translation of microcircuit-level alterations, such as the effect of NMDA receptor hyperfunction, into emergent mesoscopic population dynamics that does not explicitly model neurotransmitters dynamics. By providing a computational framework generalizing across circuit variants, Auto-MFM can thus contribute to the systematic exploration of disease-related hypotheses, enabling quantitative comparisons between in silico predictions and in vivo recordings in a broad range of brain disorders.

### 4.3 Study considerations and conclusions

Auto-MFM can be extended to support a broader range of neuronal models and new SNNs with improved biological fidelity. As an example, the phenomenological threshold in AdEx models can be represented by a second-order polynomial rather than a first-order one, requiring minimal computational modifications^22^. The generalization of the parameter-transfer strategies, including the use of PLV as a synthetic measure to transfer SNN single neuron resolution into MFM parameters, should also be validated across neuron models different from E-GLIF. It is worth noting that Auto-MFM is neither point neuron model specific tool nor a region-specific tool. It can be readily extended to other brain regions for which biologically grounded spiking models are available and have been shown to faithfully reproduce microcircuit activity, such as models of the basal ganglia and the motor cortex. This generality further supports Auto-MFM as a unifying tool for the systematic derivation of mean-field descriptions across heterogeneous brain systems.

Moreover, Auto-MFM facilitates the integration of detailed microcircuit representations into large-scale brain models, whose functional units are the MFMs^60,61^. New SNNs, with improvement in biological realism of microcircuit, are under development and MFM must follow this trend of innovation. As an example, SNNs replacing alpha-function synapses with Tsodyks–Markram dynamic synapses^62^ will require corresponding alignment in the mean field domain and Auto-MFM can substantially accelerate the derivation of functionally equivalent MFMs from updated SNN descriptions, supporting scalable mesoscale modeling while maintaining physiological interpretability, a key requirement for whole-brain simulation frameworks. By removing the need for prior expertise in MFM development, Auto-MFM systematizes the generation of biophysically-grounded population models and enables their use across research and clinical settings.

Auto-MFM (https://github.com/RobertaMLo/Auto-MFM), together with the BSB (https://www.ebrains.eu/tools/bsb)^28^ for biophysical SNN construction and TVB (https://ebrains.eu/data-tools-services/tools/the-virtual-brain)^52,53^ for whole-brain dynamics simulations, now becomes one of the building blocks of an integrated ecosystem (see Fig. 1A) that makes it possible to link macroscopic signals to their underlying microscopic causes, an otherwise inaccessible level of explanation in human in vivo studies^7^.

Crucially, it supports the incorporation of localized pathological variants, allowing the construction of virtual brains informed by synaptic-level alterations and enabling the assessment of their impact on large-scale signals. This capability provides a principled route to deconvolve macroscopic activity into its underlying cellular causes: signals that are typically linked to symptoms, such as beta-band abnormalities in Parkinson’s disease, can now be mechanistically related to specific synaptic changes. In doing so, Auto-MFM establishes a computational bridge between signals, and local circuit pathology, advancing the integration of detailed circuit knowledge into whole-brain modeling.

## Acknowledgments

This research has received funding from the European Union’s Research and Innovation Program Horizon Europe under the grant agreement No 101137289 (Virtual Brain Twin Project) to ED, and No. 101147319 (EBRAINS 2.0 Project) to ED and CC. MDG received fundings from Project PRIN 2022 - 20228B2HN5 “cerebellarNEuromodulation in ATaxia: digital cerebellar twin to predict the MOVEmentrescue (NEAT-MOVE). CGWK received funding from UKRI (#APP91962), BRC (#BRC704/CAP/CGW), MRC (#MR/S026088/1), Ataxia UK, MS Society (#77).

## Notes

### Competing Interest Statement

The authors have declared no competing interest.

