## Supplementary Material for "Automated derivation of mean field models from spiking neural networks for the simulation of brain dynamics"

### Supplementary Materials

#### MFM derivation

The cerebellar biophysical SNN is constructed using the E-GLIF model described in Geminiani et al. (2018, 2019). Single neuron parameters are reported in Tab. S1, while synaptic parameters in Tab. S2. Parameters highlighted in **bold** differ from the reference configuration, as they were specifically tuned to reproduce physiologically plausible activity of the cerebellar microcircuit.
Pareto front of TF optimization is reported in Fig.S1, and examples of the cerebellar MFM activity is reported in Fig. S2 for different input frequency.

| Parameter | Name | unit | GrC | GoC | MLI | PC |
|---|---|---|---|---|---|---|
| $g_L$ | Leak conductance | nS | 0.29 | 3.30 | 1.60 | 7.10 |
| $C_m$ | Membrane capacitance | pF | 7.00 | 145.00 | 14.60 | 334.00 |
| $\tau_{ref}$ | Refractory time | ns | 1.50 | 2.00 | 1.59 | 0.50 |
| $\tau_m$ | Membrane time constant | ns | 24.15 | 44.00 | 9.12 | 47.00 |
| $E_L$ | Resting potential | mV | -62.00 | -62.00 | -68.00 | -59.00 |
| $V_{th}$ | Threshold potential | mV | -41.00 | -55.00 | -53.00 | -43.00 |
| $V_r$ | Reset potential | mV | -70.00 | -75.00 | -78.00 | -69.00 |
| $k_{adap}$ | Adaptation constant | $MH^{-1}$ | 0.02 | 0.22 | 2.03 | 1.50 |
| k2 | Adaptation constant | $ms^{-1}$ | 0.04 | 0.02 | 1.10 | 0.04 |
| k1 | Decay rate | $ms^{-1}$ | 0.31 | 0.03 | 1.89 | 0.19 |
| A2 | Update constant | pA | -0.94 | 170.01 | 5.86 | 172.62 |
| A1 | Update constant | pA | 0.01 | 259.99 | 5.95 | 157.62 |
| $I_e$ | Endogenous current | pA | -0.89 | 16.21 | 4.45 | 700.00 |

**Tab S1. Single cell parameters for E-GLIF of the cerebellar neurons (Geminiani et al., 2018, 2019)**
Parameters specific of the type of neurons included in the multi-layer MF populations. The parameters in the top part are chosen according to literature, while the parameters at the bottom were extracted from spiking neural network simulating the cerebellar cortex spiking activity. mf = mossy fibers, GrC = Granule Cells, GoC = Golgi Cells, MLI = Molecular Layer Interneurons (Basket cells and Stellate cells)

| Pre → post synaptic | K | Q (nS) | τ (ms) |
|---|---|---|---|
| glomerulus → granule | 4 | 0.23 | **5.80** |
| glomerulus → golgi | 110 | 0.24 | **0.23** |
| golgi → granule | 5.25 | 0.24 | **13.61** |
| golgi → golgi | 3200 | 0.007 | **10.00** |
| ascending axon → golgi | 310 | 0.82 | **0.50** |
| ascending axon → purkinje | 92 | 0.41 | **1.10** |
| parallel fiber → golgi | 1100 | 0.054 | 1.25 |
| parallel fiber → purkinje | 1500 | 0.14 | **1.10** |
| parallel fiber → stellate | 520 | **0.08** | 0.64 |
| parallel fiber → basket | 840 | **0.06** | 0.64 |
| stellate → purkinje | 45.12 | 0.17 | **2.80** |
| basket → purkinje | 12 | 0.80 | **2.80** |
| stellate → stellate | 1400 | 0.005 | 2.00 |
| basket → basket | 1881 | 0.006 | 2.00 |

**Tab S2. Synaptic parameters used in the cerebellar SNN.**
The mean synaptic convergence (K) was computed as the average number of presynaptic neurons contacting a postsynaptic neuron multiplied for the number of synapses, based on the spatial connectivity rules in the

Brain Scaffold Builder (BSB). These parameters provide the microscopic reference for the mesoscale reparameterization in the MFM. For SNN-MFM parameters transfer, parameter K is multiplied for the Phase Locking Value (PLV) in parameter transfer operation as detailed in section 2.1. The quantal synaptic conductance (Q) and synaptic decay time constant (τ) were taken directly from the conductance-based synapse models implemented in the SNN. Stellate and basket cells are merged in MLI population by averaging their synaptic parameters. The parameters of GrC axonal bifurcations (parallel fiber and ascending axons) converging to the same postsynaptic population are merged parameter: K are summed, Q and τ with are averaged (see equation 13 for an example).

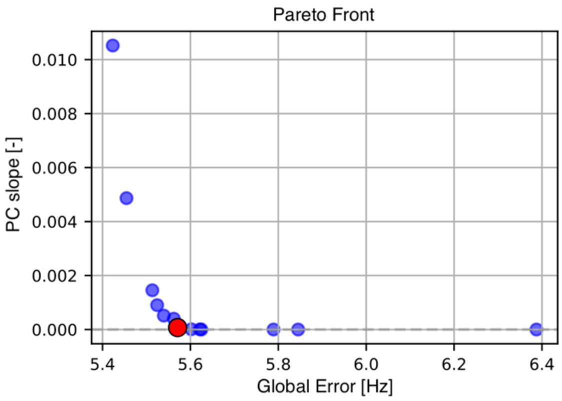


**Fig. S1) Pareto front** for α values optimization. In red the best solution identified by knee algorithm corresponding to $\alpha_{GrC}$ = 2.1, $\alpha_{GoC}$ = 2.4, $\alpha_{MLI}$ = 1.6, and $\alpha_{PC}$ = 5.4

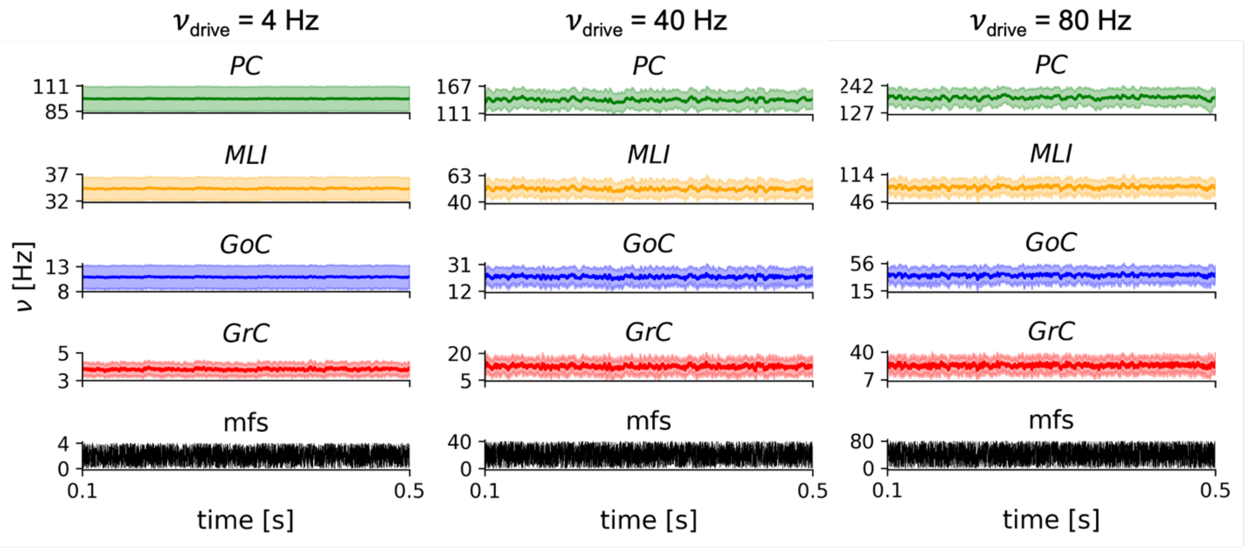


**Fig. S2) Activity of the cerebellar MFM.** Example of the MFM simulated activity for background noise like activity ($\nu_{drive}$= 4Hz, carried by mossy fibers – mfs), and mid-high input frequencies ($\nu_{drive}$= 40 Hz and $\nu_{drive}$ = 80 Hz respectively).

### Complete Taxonomy of GrC TFs

Hypo excitability configurations are derived with a reduction of the quantal synaptic parameters between mossy fibers (mfs) and GrC = $Q_{mf\text{-}GrC}$, while hyperexcitability with an increase of $Q_{mf\text{-}GrC}$. Parameter sweep is performed by spanning the values Q +- %Q. The values are reported in Tab. S3. Fig. S2 shows the numerical template, while Fig. S3 shows the analytical TFs, control configuration in the yellow box.

| % of variation | $Q_{mf\text{-}GrC}$ [nS] |
|---|---|
| -100% | 0.00 |
| -90% | 0.02 |
| -80% | 0.05 |
| -70% | 0.07 |
| -60% | 0.09 |
| -50% | 0.12 |
| -40% | 0.14 |
| -30% | 0.16 |
| -20% | 0.18 |
| -10% | 0.21 |
| | **0.23** |
| +10% | 0.25 |
| +20% | 0.28 |
| +30% | 0.30 |
| +40% | 0.32 |
| +50% | 0.35 |
| +60% | 0.37 |
| +70% | 0.39 |
| +80% | 0.41 |
| +90% | 0.44 |
| +100% | 0.46 |

**Tab S3. Qmf-GrC parametes sweep** used to derive the GrC TF taxonomy.

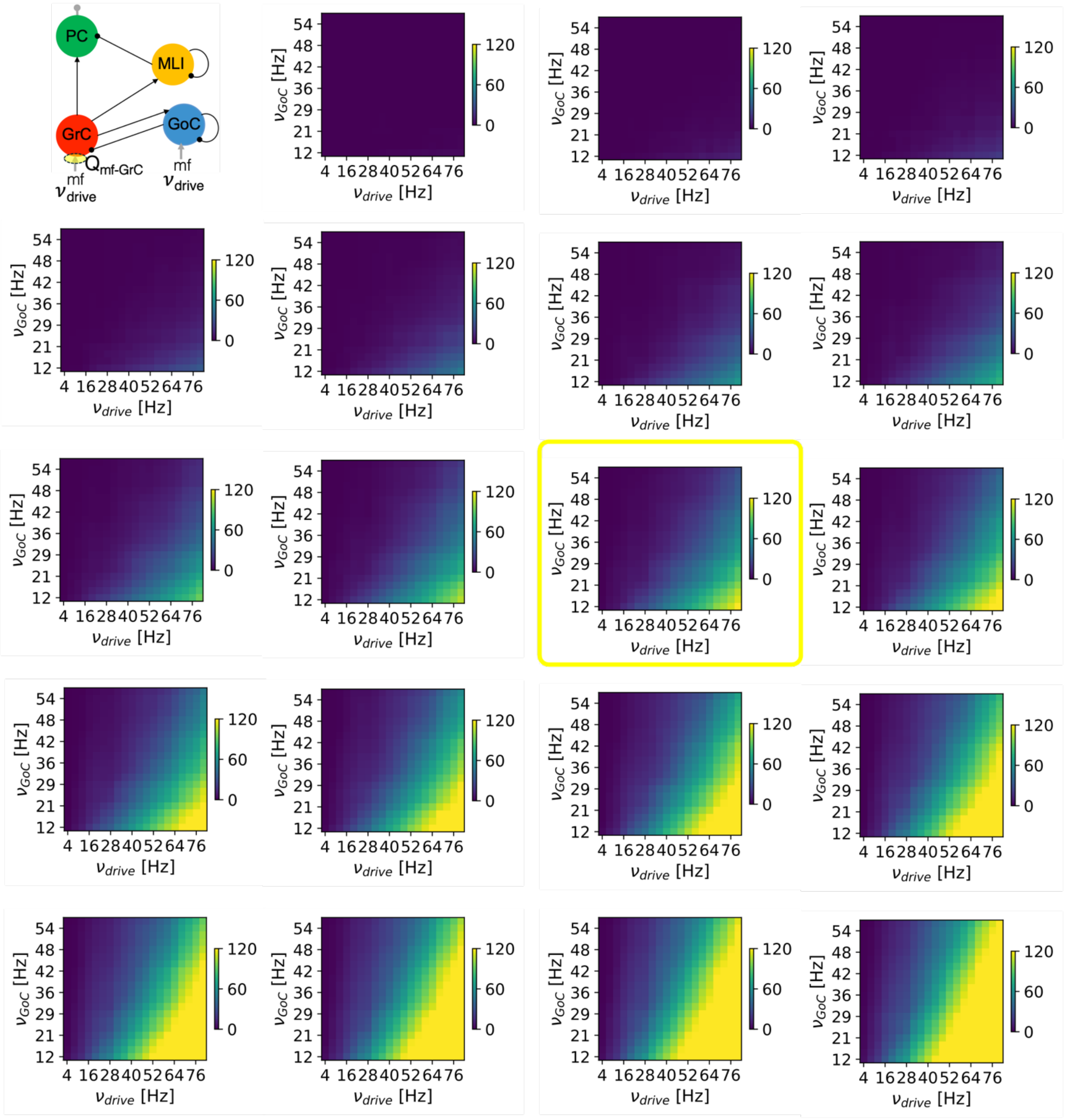


**Fig. S3) GrC TF taxonomy: numerical template.** From top left to bottom right, the parameter values increase according to the table (top left: $Q_{mf\text{-}GrC}$ = 0.0 nS; bottom right: $Q_{mf\text{-}GrC}$ = 0.46 nS). The yellow square indicates the control configuration.

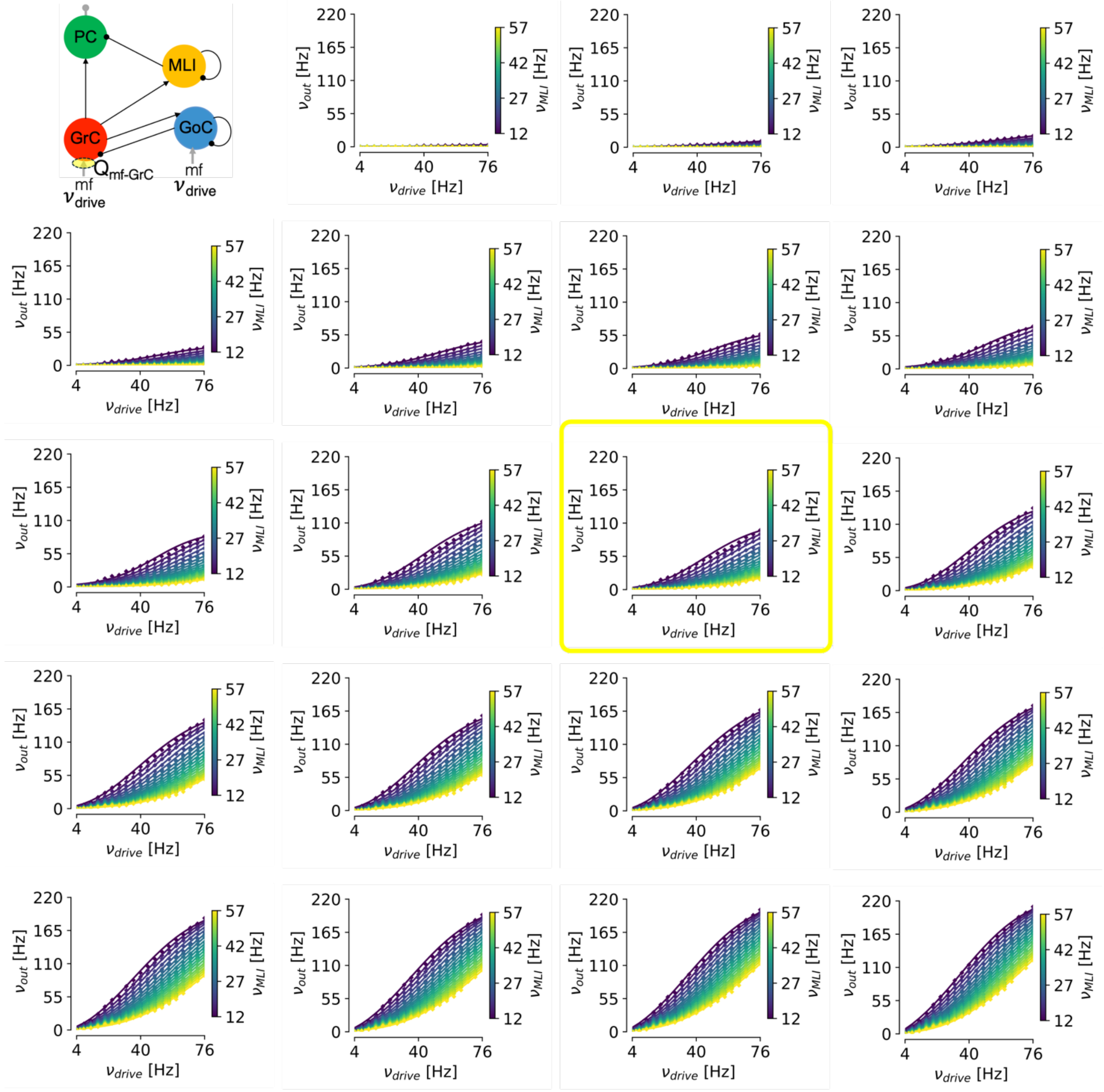


**Fig. S4) GrC TF taxonomy: analytical transfer functions,** fitted on the numerical template in Fig S2, using the two-step fitting procedure. From top left to bottom right, the parameter values increase according to the table (top left: $Q_{mf\text{-}GrC}$ = 0.0 nS; bottom right: $Q_{mf\text{-}GrC}$ = 0.46 nS). The yellow square indicates the control configuration
